# *TCF4* trinucleotide repeat expansions and UV irradiation increase susceptibility to ferroptosis in Fuchs endothelial corneal dystrophy

**DOI:** 10.1101/2022.06.27.497862

**Authors:** Sanjib Saha, Jessica M. Skeie, Gregory A. Schmidt, Tim Eggleston, Hanna Shevalye, Christopher S. Sales, Pornpoj Phruttiwanichakun, Apurva Dusane, Matthew G. Field, Tommy A. Rinkoski, Michael P. Fautsch, Keith H. Baratz, Madhuparna Roy, Albert S. Jun, Aliasger K. Salem, Mark A. Greiner

## Abstract

Fuchs endothelial corneal dystrophy (FECD), the leading indication for corneal transplantation in the U.S., causes loss of corneal endothelial cells (CECs) and corneal edema leading to vision loss. FECD pathogenesis is linked to impaired response to oxidative stress and environmental ultraviolet A (UVA) exposure. Although UVA is known to cause nonapoptotic oxidative cell death resulting from iron-mediated lipid peroxidation, ferroptosis has not been characterized in FECD. We investigated the roles of genetic background and UVA exposure in causing CEC degeneration in FECD. Using ungenotyped FECD patient surgical samples, we found increased levels of cytosolic ferrous iron (Fe^2+^) and lipid peroxidation in end-stage diseased tissues compared with healthy controls. Using immortalized and primary cell cultures modeling the *TCF4* intronic trinucleotide repeat expansion genotype, we found altered gene and protein expression involved in ferroptosis compared to controls including elevated levels of Fe^2+^, basal lipid peroxidation, and the ferroptosis-specific marker transferrin receptor 1. Increased cytosolic Fe^2+^ levels were detected after physiologically relevant doses of UVA exposure, indicating a role for ferroptosis in FECD disease progression. Cultured cells were more prone to ferroptosis induced by RSL3 and UVA than controls, indicating ferroptosis susceptibility is increased by both FECD genetic background and UVA. Finally, cell death was preventable after RSL3 induced ferroptosis using solubilized ubiquinol, indicating a role for anti-ferroptosis therapies in FECD. This investigation demonstrates that genetic background and UVA exposure contribute to iron-mediated lipid peroxidation and cell death in FECD, and provides the basis for future investigations of ferroptosis-mediated disease progression in FECD.

## INTRODUCTION

Fuchs endothelial corneal dystrophy (FECD) is a complex age-related polygenic disease that affects roughly 6.1 million Americans (1) and represents the leading indication for corneal transplant surgery in the U.S. (2-6). The condition can be diagnosed early in early stages and well before it causes visual dysfunction, through the identification of degenerative extracellular matrix deposits (guttae) on the corneal endothelium that lines the inner cornea. Gradual progression of disease results in loss of corneal endothelial cells (CECs), which do not regenerate (7), and failure to maintain appropriate corneal hydration through active ion pumping to counter the passive leakage of aqueous humor (8). Unfortunately, no available medical therapy can yet prevent disease progression, so advanced FECD requires transplantation of the endothelial cell layer to restore vision. On the molecular level, affected CECs have an increased steady-state level of reactive oxygen species (ROS), impaired antioxidant response to oxidative stress, and increased sensitivity to known exogenous stressors that drive disease progression including ultraviolet light (UV) (9-12). The cornea is particularly susceptible to damage by ultraviolet A light (UVA, 320 to 400 nm), which comprises the vast majority of incident solar radiation absorbed by CECs (9, 10, 13). Unlike ultraviolet B (UVB) light that causes DNA damage directly, UVA light causes macromolecular damage indirectly via the production of ROS that results from irradiation (14). DNA damage and apoptotic cell death in particular have been the focus of studies examining UV mediated cell death in FECD (9, 15-17). However, UVA is also known to result in lipid peroxidation and nonapoptotic oxidative cell death (18, 19). The roles of genetic background and UVA exposure in causing lipid membrane damage and endothelial cell degeneration in FECD have not been explored systematically. Specifically, the background effects of FECD mutation and exposure related effects of UVA irradiation on lipid peroxidation and CEC functioning in FECD patients or *in vivo* models have not been characterized.

A key characteristic of FECD is impaired endogenous response to oxidative stress. Vulnerability to oxidative damage in FECD has been well established, and factors that increase CEC susceptibility to lipid peroxidation have been described. Previous studies have reported decreased transcription of key antioxidant defenses in FECD including glutathione S-transferase, superoxide dismutase 2, aldehyde dehydrogenase 3A1, heme oxygenase 1, thioredoxin reductase 1, and several peroxiredoxins including Prdx 1, which protects against lipid peroxidation (20-25). Importantly, protein levels of nuclear factor erythroid 2–related factor 2 (NRF2) – the regulator of a wide-ranging metabolic response to oxidative stress, including the cystine/glutamate antiporter (system *x_c_^-^*) that imports cysteine for glutathione biosynthesis – are reduced in FECD (20, 26, 27). Normal levels of glutathione (GSH) and normal functioning of glutathione peroxidase 4 (GPX4), which catalyzes the reduction of lipid peroxides in a GSH-dependent reaction, are important for protecting cells against nonapoptotic oxidative cell death via iron-dependent lipid peroxidation, or ferroptosis (28-30). Of note, GPX4 levels are lower than controls in FECD surgical samples, indicating that FECD increases susceptibility to ferroptosis (31, 32). A possible role for ferroptosis in FECD has been postulated but not investigated to account for the increased susceptibility to oxidative damage and lipid peroxidation (4, 33). In addition to increased ROS-mediated lipid peroxidation and decreased GPX4 functioning, altered iron metabolism is required for ferroptosis to occur (34, 35). In health, ferric iron (Fe^3+^) is bound to transferrin, imported into the cell via transferrin receptor 1 (TFR1, also known as CD71) mediated endocytosis, and stored in ferritin (36, 37). A pool of labile and biologically reactive Fe^2+^ is available, but levels are carefully regulated in the cell (37, 38). In pathological exposures to oxidative stress, an excess of free Fe^2+^ reacts with membrane-bound lipids to cause ferroptosis (34). Reactive Fe^2+^ at or near cell membranes can drive Fenton reactions, which cause the formation of toxic prooxidant radicals (unstable) and non-radical lipid hydroperoxide intermediaries (stable and detectable) and result in membrane-bound lipid peroxidation (39-43). This is of particular interest in FECD given the increased susceptibility to UVA-induced damage in affected patients. UV exposure is known to result in iron accumulation and ferroptosis (44-46), and iron release from ferritin with UVA irradiation has been well reported in literature (45, 47, 48). To date, no studies characterizing the role of iron, iron-lipid perturbations, or lipid peroxidation in FECD pathobiology have been reported.

Although multiple lines of evidence support the theoretical basis for ferroptosis to be a pathological component of cell death in FECD, ferroptosis has not been characterized or evaluated systematically in this disease. A detailed understanding of ferroptosis in FECD would facilitate the development of targeted pharmacological therapies directed at preventing oxidative damage and oxidative cell death. We hypothesized that both FECD genetic background and UVA increase CEC susceptibility to nonapoptotic oxidative cell death and lipid peroxidation through the accumulation of toxic intracellular concentrations of ferrous iron. To test this hypothesis, we studied ungenotyped FECD patient surgical samples and immortalized and primary cell culture models from FECD patients with pathological expansions of trinucleotide repeats in intron 3 of the *TCF4* gene (the most common genotype associated with FECD (49-51)) and evaluated for increases in lipid peroxidation, cytosolic Fe^2+^, and susceptibility to ferroptosis attributable to genetic background and UVA exposure. We also evaluated for the expression of key genes and proteins associated with ferroptosis, including TFR1 (responsible for the influx of iron inside the cell), FPN1 (efflux transporter of intracellular iron), FSP1 (ferroptosis suppressor protein 1, inhibits ferroptosis by NAD(P)H-dependent reduction of ubiquinone to ubiquinol), FTH (ferritin heavy chain, converts Fe^2+^ to Fe^3+^ and stores iron to maintain homeostasis), FTL (ferritin light chain, iron reservoir and removes excess iron), and GPX4 (inhibits lipid peroxidation by converting hydroperoxides into lipid alcohols) and the capacity for molecules with anti-ferroptotic activity to prevent key cellular processes implicated in ferroptosis (36, 52-56). Results of this investigation provide a basis for future mechanistic investigations of ferroptotic cell death and the prevention of disease progression in FECD.

## RESULTS

### FECD surgical samples demonstrate key markers of ferroptosis

To gain insight into whether ferroptosis plays a role in FECD pathogenesis, we first performed an analysis of published RNA-Seq datasets of samples from 36 FECD and 8 control patients for the presence of genes known to be involved in ferroptosis (57). Gene set enrichment analysis showed downregulation of a ferroptotic gene signature in FECD patients compared to control patients (FDR < 0.01); however, a mixed model analysis that allows for both upregulated and downregulated genes was even more significantly enriched (FDR < 0.001, **Fig. 1A**), implicating the involvement of ferroptosis in FECD pathogenesis.

**Figure 1:**
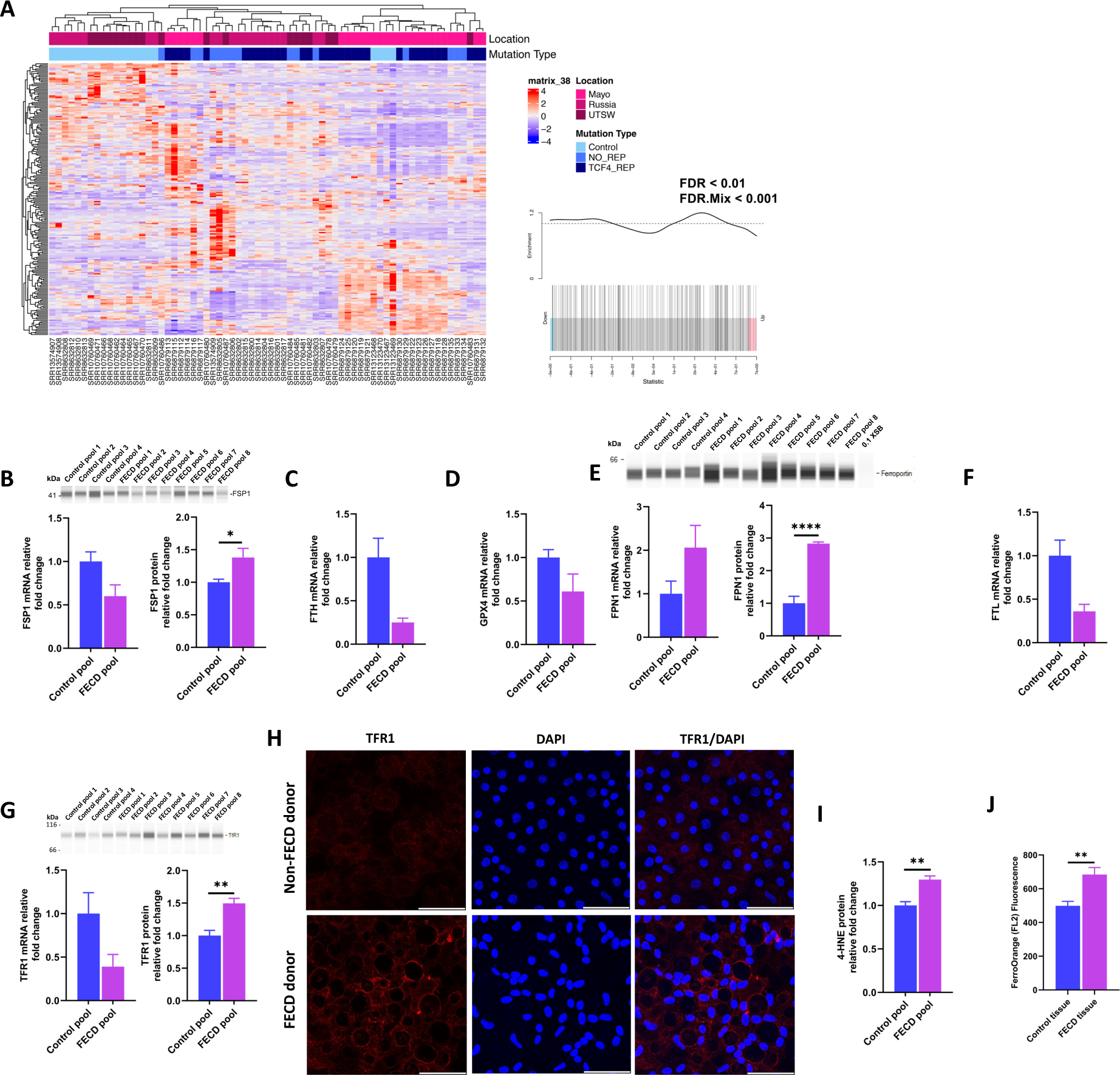
FECD surgical tissues show key markers of ferroptosis. (**A**) Heatmap with hierarchical clustering for 211 genes from the FerrDB database that includes known driver, suppressor, and marker ferroptosis genes that were expressed in the RNA-Seq datasets. For each plot, “pearson” was used for the clustering distance and “complete” for the hierarchical clustering method. The location where the representative dataset was collected (Mayo, Russia, or UTSW) and mutation type (Control, no TCF4 repeats [No_Rep] or TCF repeats [TCF4_Rep]) are shown for each sample. (**B**) *FSP1* mRNA and protein expression in FECD and control tissues. **(C)** *FTH* mRNA expression in control and FECD tissues. **(D)** *GPX4* mRNA expression in FECD and control human tissues. **(E)** Ferroportin (*FPN1*) mRNA and protein expression in FECD and control tissues. (**F**) *FTL* mRNA expression in control and FECD tissues. (**G**) TFR1 mRNA and protein expression in control and FECD surgical tissues. (**H**) Representative immunohistochemistry images of TFR1 localization in non-FECD and FECD donor cornea tissues. **(I)** 4-HNE protein expression in human surgical samples from patients with FECD (n = 8). All data of mRNA and protein expression are shown as mean±SEM for n = 12 (Control tissues, 8 pools of 3, each pool contained 3 tissues) and n = 24 (FECD tissues, 8 pools of 3, each pool contained 3 tissues). All the statistical comparisons were conducted using two-tailed, unpaired Student’s t-test, *p < 0.05, **p < 0.01, ****p < 0.0001. Relative gene expression is normalized by *β-actin*. **(J)** Cytosolic Fe^2+^ in primary CECs isolated from healthy human donor corneas (n = 11, each cornea divided into 2 sections) and FECD surgical explants (n = 7). Data are shown as mean±SEM; **p < 0.01, Student’s t-test.

Next, we sought to validate these findings in our own corneal endothelial cell-Descemet membrane tissue samples resected from ungenotyped FECD patients (**Supplementary table 1 and 2**) by evaluating for expression of key genes and proteins associated with ferroptosis, including FPN1, FSP1, FTH, FTL, GPX4 and TFR1 (35, 58, 59). Consistent with the published RNA-Seq FECD datasets, our FECD surgical samples showed that *FSP1, FTH*, and *GPX4* gene expression was decreased by 0.6, 0.25, and 0.62-fold, respectively (**Fig. 1B-D**), and *FPN1* gene expression was increased by 2.06-fold in (**Fig. 1E**). *FTL* and *TFR1* gene expression was downregulated by 0.36 and 0.39-fold in our samples (**Fig. 1F-G**) but showed no difference in the published datasets. Consistent with the gene expression data, FPN1 showed a 2.83-fold increase in protein expression (p < 0.001, **Fig. 1E**), indicating a compensatory response by FECD to export higher labile iron in order to restore iron homeostasis. Interestingly, though *FSP1* and *TFR1* showed decreased gene expression, protein expression in our surgical samples was significantly upregulated by 1.38 and 1.50-fold, respectively (p < 0.05 and p < 0.01, respectively) (**Fig. 1B and 1G**). *TFR1*, which internalizes transferrin-iron complexes through endocytosis, serves as a useful and specific marker of ferroptosis because it contributes to higher iron intake and correlates with iron-mediated lipid peroxidation (36). Notably, immunohistochemistry images show that TFR1 protein expression was higher and localized more to the surface of CECs in FECD donor tissues than non-FECD donor tissues (**Fig. 1H**).

In addition to key ferroptotic expression markers, we assessed whether key cellular processes implicated in ferroptosis, such as lipid peroxidation and accumulation of cytosolic ferrous iron [Fe^2+^], were increased in our FECD surgical samples. Accordingly, FECD surgical samples showed 1.3-fold greater accumulation of lipid peroxidation end products compared to control donor tissues (p < 0.01) (**Fig. 1I, Supplementary figure 1**) and 32% more cytosolic Fe^2+^ content in isolated CECs (p < 0.01) (**Fig. 1J, Supplementary table 3**).

Furthermore, we collected aqueous humor samples from 4 patients with and 4 patients without FECD (**Supplementary table 4**) and conducted protein mass spectrometry analysis to look for further evidence of iron dysregulation. A total of 23,171 protein isoforms were identified, of which 3,448 protein isoforms had significantly altered expression in FECD patient aqueous compared to non-FECD aqueous samples (**Supplementary table 5-6**) (60). Ingenuity molecular pathway analysis of the statistically significant proteins (p < 0.05) between FECD and control aqueous humor identified ferroptosis as one of the top 15 most significantly represented pathways. Several isoforms of human transferrin were higher in FECD aqueous samples compared to controls (3.8 < Ratio < 18.7; p < 0.015), consistent with previous findings that receptor-mediated endocytosis of the transferrin-Fe^3+^ complex is required for ferroptosis (61). A complete list of differentially expressed proteins can be found in **supplementary table 6**.

### Increased oxidative stress, lipid peroxidation, and iron overload in FECD cells

Due to the scarcity of surgical FECD tissue samples, we utilized an established *TCF4* expanded repeat CEC line derived from an FECD patient (F35T), human *TCF4* expanded repeat primary CEC lines from FECD patients, and a control CEC line (HCEC-B4G12) for additional experiments. As oxidative stress is highly implicated in FECD pathogenesis, we first recapitulated our and others work (3, 10, 12, 33, 62) by showing that *TCF4* expanded repeat F35T cells had a higher basal ROS (p < 0.0001, **Fig. 2A-B**) and basal mitochondrial superoxide concentration (p < 0.0001, **Fig. 2C**) than control HCEC-B4G12 cells. Consistent with our findings from surgical samples, *TCF4* expanded repeat F35T cells had 0.47-fold lower GPX4 expression (p < 0.001; **Fig. 2D)** than control HCEC-B4G12 cells and a commensurate 2.89-fold greater basal level of lipid peroxidation (p < 0.0001; **Fig. 2E-F**). *GPX4* gene expression was upregulated by 2.27-fold in F35T cells when compared to B4G12 cells though protein was downregulated (**Fig. 2D**). Primary cells from FECD patients with *TCF4* expanded repeats had 1.60-fold higher GPX4 expression than non-FECD primary cells (**Fig. 2G**).

**Figure 2:**
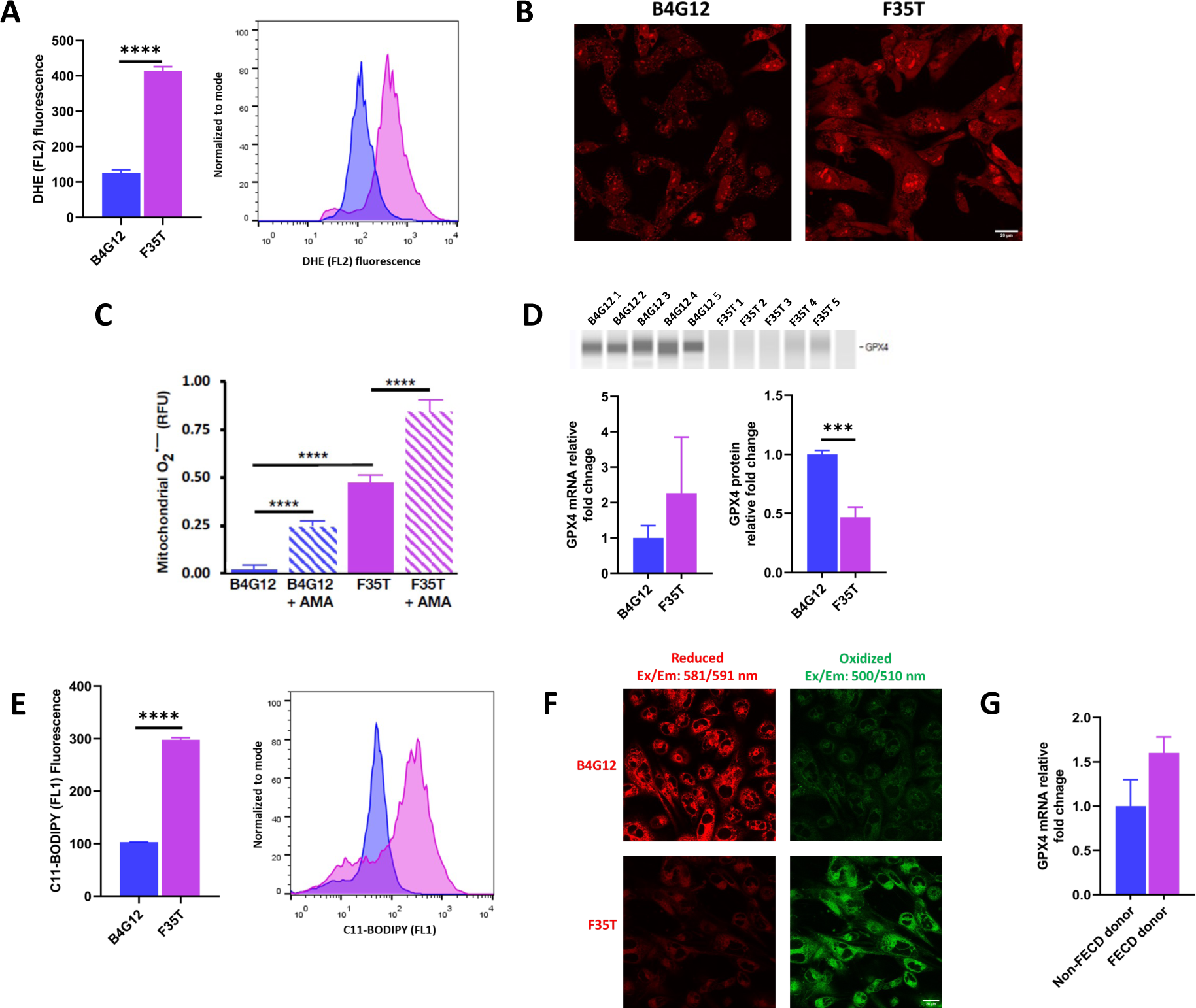
FECD immortalized and primary cell culture shows key marker of ferroptosis. **(A)** Representation of median of the fluorescence of DHE showing significant difference in ROS between indicated cells. DHE (FL2 fluorescence) peak of F35T cells shifts to right when compared to B4G12 cells. **(B)** Representative confocal images showing fluorescence of DHE indicating ROS in the indicated cell lines. **(C)** Mitochondrial ROS quantified by MitoROS 580 dye in the indicated cells. Data are shown as mean±SEM; n = 3; ****p < 0.0001, one-way ANOVA, followed by Tukey’s post-hoc test. AMA indicates antimycin-A**. (D)** *GPX4* mRNA and protein expression in HCEC-B4G12 and F35T cells. (**E**) Basal level of lipid peroxidation in HCEC-B4G12 and F35T cells quantified by C11-BODIPY fluorescent probe using flow cytometry. Comparisons of median fluorescence of C11-BODIPY detected in HCEC-B4G12 and F35T cells (10,000 cells). Data are shown as mean±SEM; n = 3; ****p < 0.0001, Student’s t-test. C11-BODIPY (FL1 fluorescence) peak of F35T cells shifts to right when compares to B4G12 cells. (**F**) Representative confocal images showing fluorescence of reduced and oxidized dye in the indicated cell lines. (**G**) *GPX4* mRNA expression in human expanded *TCF4* repeat expansion primary cells. All data of mRNA and protein expression are shown as mean±SEM for n = 5-9 (B4G12), n = 5-7 (F35T) and n = 4 (both non-FECD and FECD donor primary cells). All the statistical comparisons were conducted using two-tailed, unpaired Student’s t-test, ***p < 0.001. Relative gene expression is normalized by *β*-actin.

In addition to GPX4 dysregulation and ROS-mediated lipid peroxidation, altered iron metabolism and disruption of iron homeostasis leading to toxic concentrations of labile intracellular Fe^2+^ is required for ferroptotic cell death (34, 35). Consistent with this, *TCF4* expanded repeat F35T cells showed a 3.3-fold greater cytosolic (p < 0.0001) and 10.9-fold greater mitochondrial (p < 0.0001) Fe^2+^ accumulation than control HCEC-B4G12 cells (**Fig. 3A-F**), which was associated with a 5.4-fold upregulation of TFR1 (p < 0.001, **Fig. 3G-H)** and a 20% and 72% decrease in total ferritin (p < 0.001, **Fig. 3I**) and mitochondrial ferritin (FtMt) protein (p < 0.0001, **Fig. 3J**), respectively. *TCF4* expanded repeat F35T cells showed upregulation of *TFR1*, *FTH* and *FTL* mRNA by 4.77, 2.37 and 2.64-fold, respectively, in comparison to control cells (**Fig. 3H and 3K-L**). *FSP1* mRNA and FSP1 protein were upregulated by 2.51 and 1.61-fold in F35T cells, respectively (**Fig. 3M**). In agreement with findings in F35T cells, *TFR1*, *FTH*, *FTL* and *FSP1* mRNA were upregulated in primary cells from FECD patients with *TCF4* expanded repeats (**Fig. 3N-Q**). The contrast between tissues and cells may reflect changes attributable to disease stage and/or model differences. Of note, ferritin can store labile Fe^2+^ inside its multimeric shell in a nontoxic state (63) and FtMt can safely redistribute toxic iron from the cytosol to the mitochondria and protect mitochondria from iron-mediated oxidative damage (64-66). However, the presence of increased intracellular Fe^2+^ in the context of decreased ferritin and FtMt indicates intracellular overloading of toxic Fe^2+^. Moreover, iron overload due to both higher iron uptake associated with TFR1 overexpression and downregulation of ferritin protein via ferritinophagy, which further increases TFR1 expression, have both been reported to lead to accumulation of toxic Fe^2+^ and promote ferroptosis (63).

**Figure 3:**
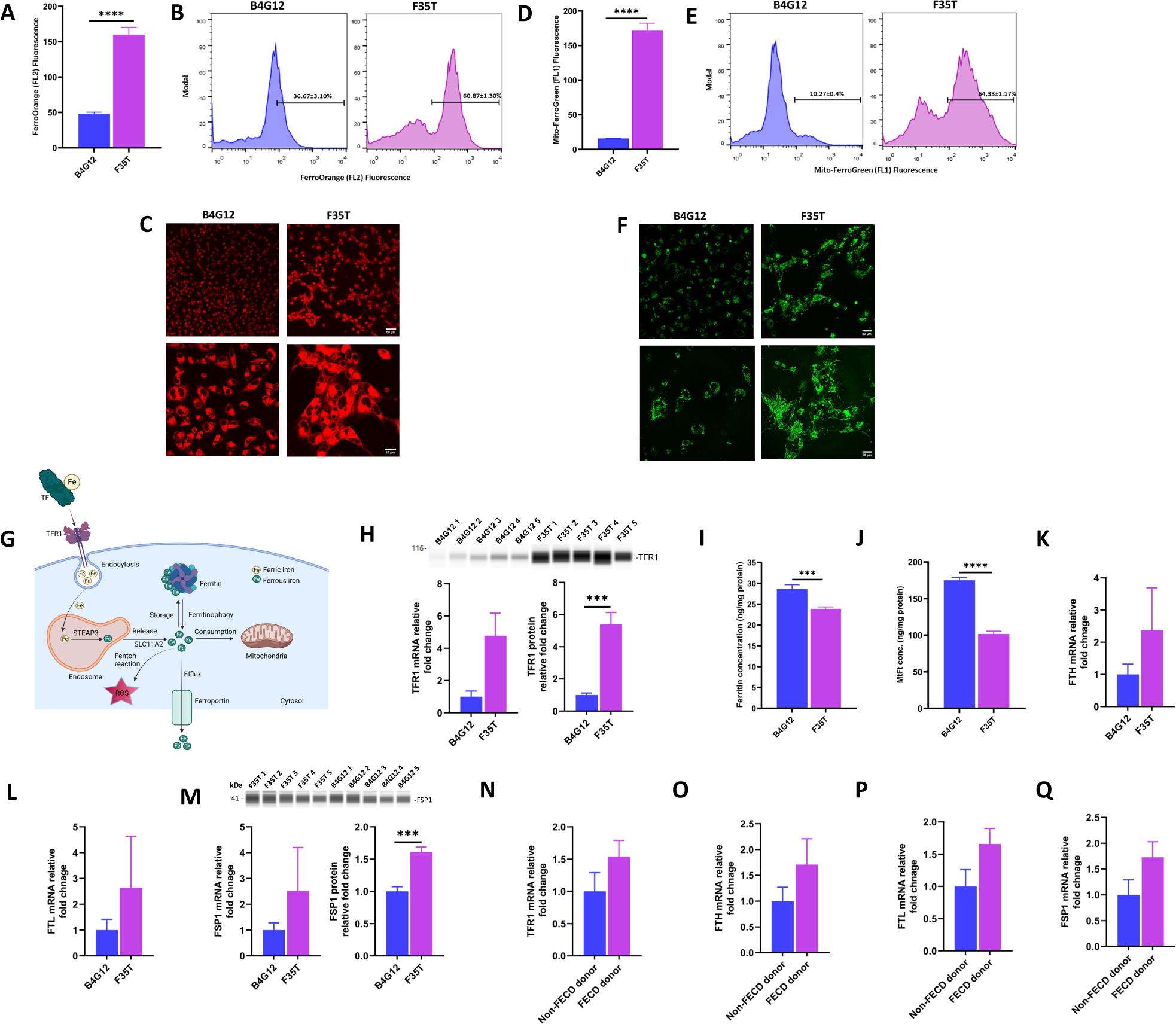
FECD immortalized and primary cell culture show key gene and protein expression changes related to ferroptosis. (**A**) Representation of median of the fluorescence of FerroOrange showing significant difference in cytosolic Fe^2+^ between indicated cells. B4G12 and F35T cells were stained with FerroOrange fluorescent probe to quantify cytosolic Fe^2+^ by flow cytometry. A minimum 10,000 cells were quantified for measuring the fluorescence. **(B)** Cell population distribution after staining with FerroOrange showing cytosolic Fe^2+^ content. **(C)** Representation of confocal microscopy images of HCEC-B4G12 and F35T cells stained with FerroOrange fluorescent probe showing and comparing cytosolic Fe^2+^. **(D)** Representation of median of the fluorescence of Mito-FerroGreen showing significant difference in mitochondrial Fe^2+^ between indicated cells. HCEC-B4G12 and F35T cells were stained with Mito-FerroGreen fluorescent probe to quantify mitochondrial Fe^2+^ by flow cytometry. A minimum of 10,000 cells were quantified for measuring fluorescence. **(E)** Cell population distribution after staining with Mito-FerroGreen showing mitochondrial Fe^2+^ content. **(F)** Representation of confocal microscopy images of HCEC-B4G12 and F35T cells stained with Mito-FerroGreen fluorescent probe showing and comparing mitochondrial Fe^2+^. Data are represented as means ± SEM; n = 3; unpaired Student’s t-test, ****p < 0.0001. **(G)** Representation of cellular iron metabolism in ferroptosis process. (**H**) *TFR1* mRNA and protein expression in HCEC-B4G12 and F35T cells. **(I)** Quantification of human ferritin (Ft) by ELISA in protein from HCEC-B4G12 and F35T cells. The human ferritin levels are presented as mean±SEM; n = 9 (3 biological replicates × 3 technical replicates); ***p <0.001, Student’s t-test. (**J)** Quantification of mitochondrial ferritin (FtMt) by ELISA in protein from HCEC-B4G12 and F35T cells. The mitochondrial ferritin levels are presented as mean±SEM; n = 9 (3 biological replicates × 3 technical replicates); ****p < 0.0001, Student’s t-test. **(K)** *FTH* mRNA expression in indicated cells. **(L)** *FTL* mRNA expression in indicated cells. (**M**) *FSP1* mRNA and protein expression in the indicated cells. (**N**) *TFR1* mRNA expression in non-FECD and FECD donor expanded *TCF4* repeat expansion primary cells. **(O)** *FTH* mRNA expression in non-FECD and FECD donor expanded *TCF4* repeat expansion primary cells. **(P)** *FTL* mRNA expression in non-FECD and FECD donor primary cells. (**Q**) *FSP1* mRNA expression in primary cells. All data of mRNA and protein expression are shown as mean±SEM for n = 5-9 (B4G12), n = 5-7 (F35T) and n = 4 (primary cells). All the statistical comparisons were conducted using two-tailed, unpaired Student’s t-test, ***p < 0.001, ****p < 0.0001. Relative gene expression is normalized by *β-actin*.

### Increased susceptibility to RSL3-induced ferroptosis in FECD cells

Increased ROS and lipid peroxidation in the setting of iron imbalance are the key hallmarks of ferroptosis; therefore, we evaluated whether further cellular perturbation with RSL3, an inhibitor of GPX4 peroxidase activity and well known inducer of ferroptosis (67), would promote greater than expected ferroptotic cell death in F35T cells compared to HCEC-B4G12 control cells. Following RSL3-induced ferroptosis, we observed a dramatic, early increase in the number of lipid droplets in both F35T and control cells (1-3 h), with F35T cells showing more droplets at baseline (p < 0.0001, **Fig. 4A-B**) and after treatment (**Fig. 4C-D, Supplementary video 1A, 1B**) (68); this was followed by a rapid and dramatic decrease in visible droplets that occurred just prior to cell death (≥4 h) and correlated with increased lipid peroxidation (**Fig. 4D, 6B, Supplementary video 1A-B**). Taken together, we and others (68) show that lipid droplets initially form to maintain cellular energy homeostasis and prevent lipotoxicity; however, when this compensatory mechanism becomes overwhelmed, cells undergo lipophagy to induce lipid release, lipid peroxidation and subsequent ferroptotic cell death. Consequently, F35T cells showed increased ferroptosis susceptibility, undergoing ferroptotic cell death approximately 1 h earlier than control cells as evidenced by both Sytox cell death assay and light microscopy, with the latter demonstrating typical ferroptosis morphological changes including cellular swelling, presence of lipid droplets at the perinuclear space at the time of death, and plasma membrane rupture (**Fig. 4D, Supplementary video 1A-B, 2A-B**). Additionally, F35T cells demonstrated significantly more cell death than control cells (p < 0.01, **Fig. 5A-B**), reached peak levels of cell death 1.5 h sooner than control cells, and displayed a characteristic ferroptosis wave-like pattern of propagating lipid damage and cell death **(Supplementary figure 2)** (69). Of note, iron-chelation therapy with deferoxamine (DFO) did protect control cells from RSL3-induced ferroptosis between 4-8 h (p < 0.0001; **Fig. 4E**) but had minimal effect on F35T cells, providing further evidence of increased ferroptosis susceptibility in FECD cells.

**Figure 4:**
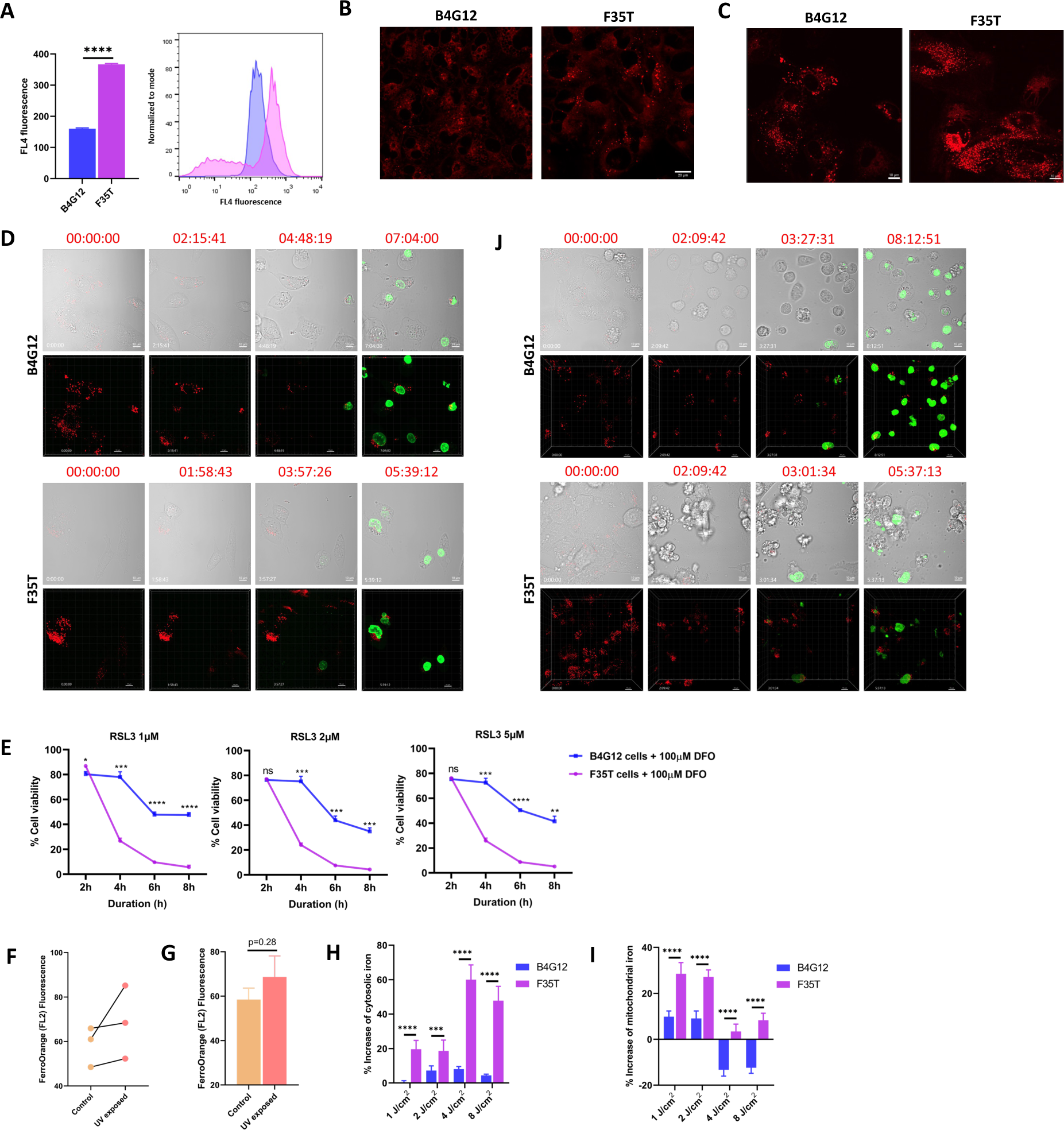
FECD demonstrates higher lipid droplets, iron overload and higher susceptibility to ferroptosis induced by RSL3 and UVA irradiation. **(A)** Representation of median fluorescence of LipidSpot™ 610 showing significant difference in ROS between indicated cell lines. LipidSpot™ 610 (FL4 fluorescence) peak of F35T cells shifts to right when compared to B4G12 cells. **(B)** Representative confocal images showing fluorescence of LipidSpot™ 610 indicating lipid droplets in the indicated cell lines. In flow cytometer analysis, data are shown as mean±SEM; n = 9 (3 biological replicates × 3 technical replicates); ****p < 0.0001, Student’s t-test. **(C)** Representative confocal images showing RSL3 induced lipid droplets in the indicated cells. (**D**) Time lapse confocal images of ferroptosis events induced by RSL3. **(E)** Deferoxamine (DFO) iron chelation assay where HCEC-B4G12 and F35T cells were treated with DFO for 24h and then challenged with RSL3 at 1, 2 and 5 µM for different durations of 2, 4, 6 and 8h. Cell viability was measured by MTS assay. Data are represented as mean ± SEM for three biological replicates; Student’s t-test, * p < 0.05, ** p < 0.01, *** p < 0.001, **** p < 0.0001. **(F)** Cytosolic Fe^2+^ release in human donor corneas upon UVA irradiation at 5 J/cm^2^. Primary endothelial cells were isolated after UVA irradiation and stained with FerroOrange fluorescent probe. The experiment was conducted pairwise, where the left cornea was used as control and the right cornea was exposed to UVA. **(G)** Comparison of geometric mean of FerroOrange (FL2 fluorescence) signals. Data are mean±SEM, Paired two-tailed Student’s t-test. **(H)** Representation of percent increase of labile cytosolic Fe^2+^ after UVA irradiation at the fluence of 1, 2, 4 and 8 J/cm^2^. Indicated cells were stained with FerroOrange fluorescent probe immediately post-UVA (Data are shown as mean±SEM, n = 9; 3 biological replicates × 3 technical replicates; ***p < 0.001, ****p < 0.0001, one-way ANOVA, followed by Tukey’s post-hoc test). **(I)** Representation of percent increase of mitochondrial Fe^2+^ after UVA irradiation at the fluence of 1, 2, 4 and 8 J/cm^2^. Indicated cells were stained with Mito-FerroGreen fluorescent probe immediately post-UVA (Data are shown as mean±SEM, n = 9; 3 biological replicates × 3 technical replicates; ****p < 0.0001, one-way ANOVA, followed by Tukey’s post-hoc test.). (**J**) Time lapse confocal images of ferroptosis events induced by UVA irradiation at 1.5 J/cm^2^. Representative confocal images showing disappearance of lipid droplets at the time of ferroptosis induced death in the indicated cells. SYTOX green indicates cell death.

**Figure 5:**
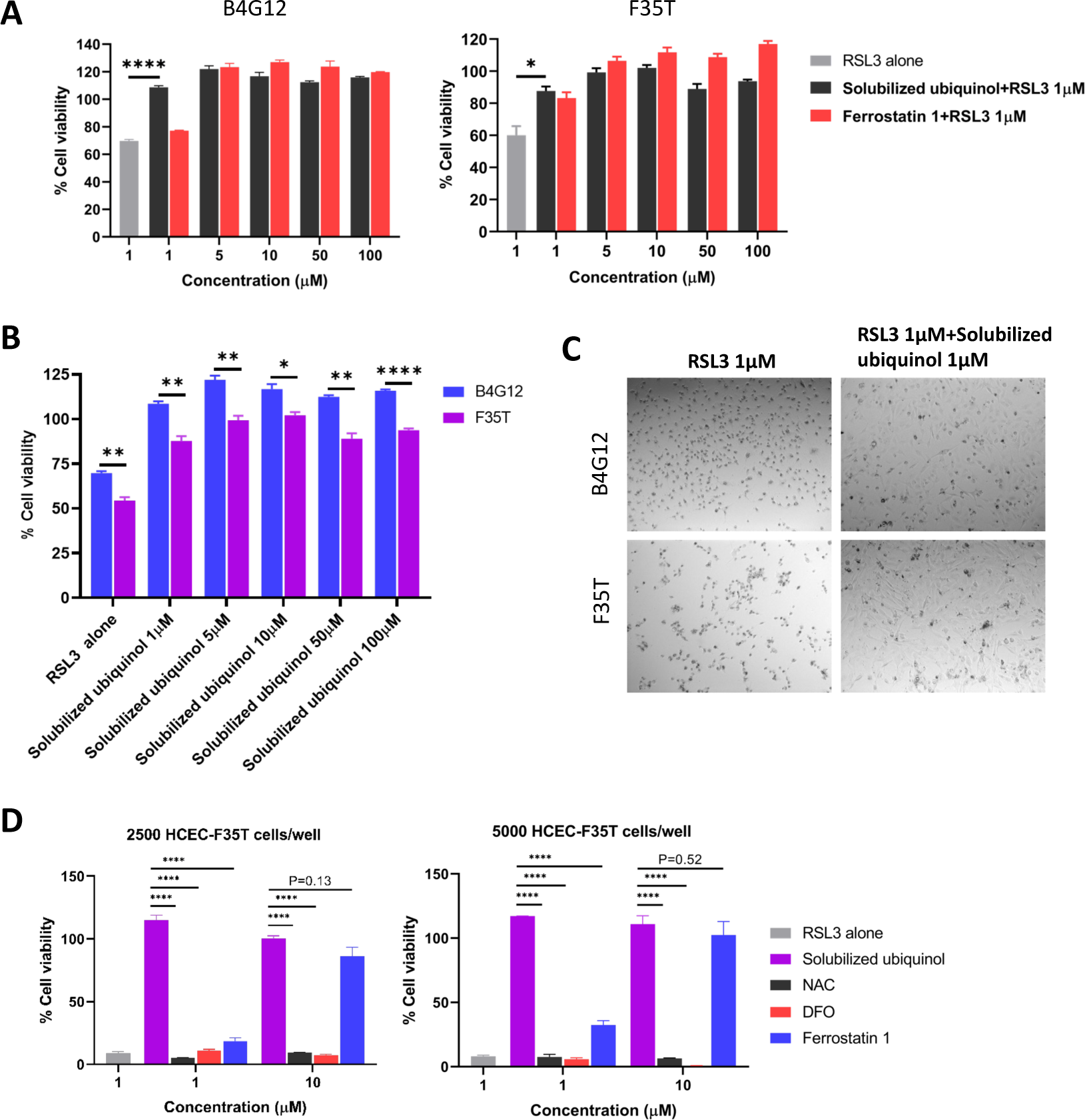
Solubilized ubiquinol gives protection against ferroptosis. **(A)** Cell viability assay was performed by quantifying LDH release after treatment with solubilized ubiquinol at different concentrations of 1, 5, 10, 50, and 100 µM. Following solubilized ubiquinol treatment, HCEC-B4G12 and F35T cells were challenged with 1 µM of RSL3 for 24 h. Data are shown as mean±SEM; n = 3; ****p < 0.0001, Student’s t-test against RSL3 alone group. **(B)** Comparisons of cell viability of HCEC-B4G12 and F35T cells following the treatment of solubilized ubiquinol. Solubilized ubiquinol was comparatively less effective in F35T cells, indicating more ferroptosis compared to B4G12 cells. Data are shown as mean±SEM; n = 3; *p < 0.05, **p < 0.01, **** p < 0.0001, Student’s t-test. **(C)** Representative light microscopy images showing that solubilized ubiquinol at 1 µM dose inhibited RSL3 induced ferroptosis in HCEC-B4G12 and F35T cells. **(D)** Solubilized ubiquinol outperforms NAC, DFO and ferrostatin-1 in protecting F35T cells against RSL3 induced cell death in ferroptosis assay. Data are shown as mean±SEM, n = 3; **** p < 0.0001, Student’s t-test.

### UVA exposure induces iron overload and ferroptosis in FECD cells

As UVA is a major component of sunlight that causes severe oxidative stress in exposed cells (70, 71), increases labile cytosolic Fe^2+^ release from the core of cytosolic and mitochondrial ferritins (45, 47, 71), and is implicated in FECD pathogenesis and progression (9, 15-17), we considered whether UVA may exert its toxic effects in susceptible FECD cells by inducing iron overload and promoting ferroptosis. To evaluate this, we first treated F35T cells with varying amounts of UVA radiation to determine a physiological dose of UVA irradiation (5 J/cm^2^) that minimally affected cell viability (**Supplementary figure 3**). Upon UVA irradiation, labile cytosolic Fe^2+^ was increased in a pair-wise comparison with control unexposed corneas (p = 0.28; **Fig. 4F-G, Supplementary table 7**), confirming that even subthreshold levels of UVA irradiation can result in CEC iron release and intracellular accumulation.

Next, we exposed both F35T and HCEC-B4G12 control cells to various doses of UVA irradiation above and below the predetermined subthreshold level (1, 2, 4 and 8 J/cm^2^). Upon irradiation, F35T cells had a significantly (p < 0.0001) higher increase in cytosolic Fe^2+^ accumulation (20-60%) when compared with control cells (0-8%) at all doses of UVA irradiation tested (**Fig. 4H**), indicating increased susceptibility of F35T cells to UVA-mediated toxicity. Similarly, F35T cells had a significantly higher percentage of mitochondrial Fe^2+^ accumulation than control cells at all doses of UVA irradiation (p <0.0001, **Fig. 4I**). At higher dose levels of 4 and 8 J/cm^2^, we observed a decrease in mitochondrial Fe^2+^ levels in both cell lines compared to lower irradiances, suggestive of UVA-induced mitochondrial fragmentation (72, 73) in which Fe^2+^ is released into the cytosol (17). Following UVA exposure of 1.5 J/cm^2^, we observed strikingly similar morphological findings in F35T cells compared to control cells consistent with RSL3-induced ferroptosis, including a dramatic increase in lipid droplets at baseline and after exposure (**Fig. 4J**), a sharp decrease in lipid droplets prior to cell death (3-4 h), earlier onset of cell death (0.5 h before control cells as evidenced by both Sytox cell death assay and light microscopy), earlier peak levels of cell death (2.5 h sooner than control cells), and typical ferroptosis morphological changes including cellular swelling, presence of lipid droplets at the perinuclear space at the time of death, and plasma membrane rupture (**Fig. 4J, Supplementary video 3A-B**).

### Ubiquinol prevents RSL3-induced ferroptosis in FECD and healthy cells

Next, in order to identify potential therapeutic options for FECD, we evaluated several pharmacological compounds known to protect against lipid peroxidation, iron overload, and ferroptosis, including solubilized ubiquinol (33), ferroptosis inhibitor ferrostatin-1, antioxidant N-acetyl cysteine (NAC), and iron-chelator deferoxamine (DFO). Strikingly, after 24 hours of RSL3 treatment, solubilized ubiquinol outperformed all other tested compounds in reducing lipid peroxidation and preventing cell death in both cell lines (**Fig. 5-6**), completely abrogating RSL3-induced ferroptotic cell death at low concentrations (1-5 µM) without any evidence of morphological cellular damage (**Fig. 5A-C**). Ferrostatin-1 provided a small but significant rescue at 1 µM in preventing cell death with complete rescue at 5-10 µM, whereas NAC and DFO had no effect at the tested conccentrations in preventing RSL3-induced ferroptosis at 24 hours (**Fig. 5D**).

**Figure 6:**
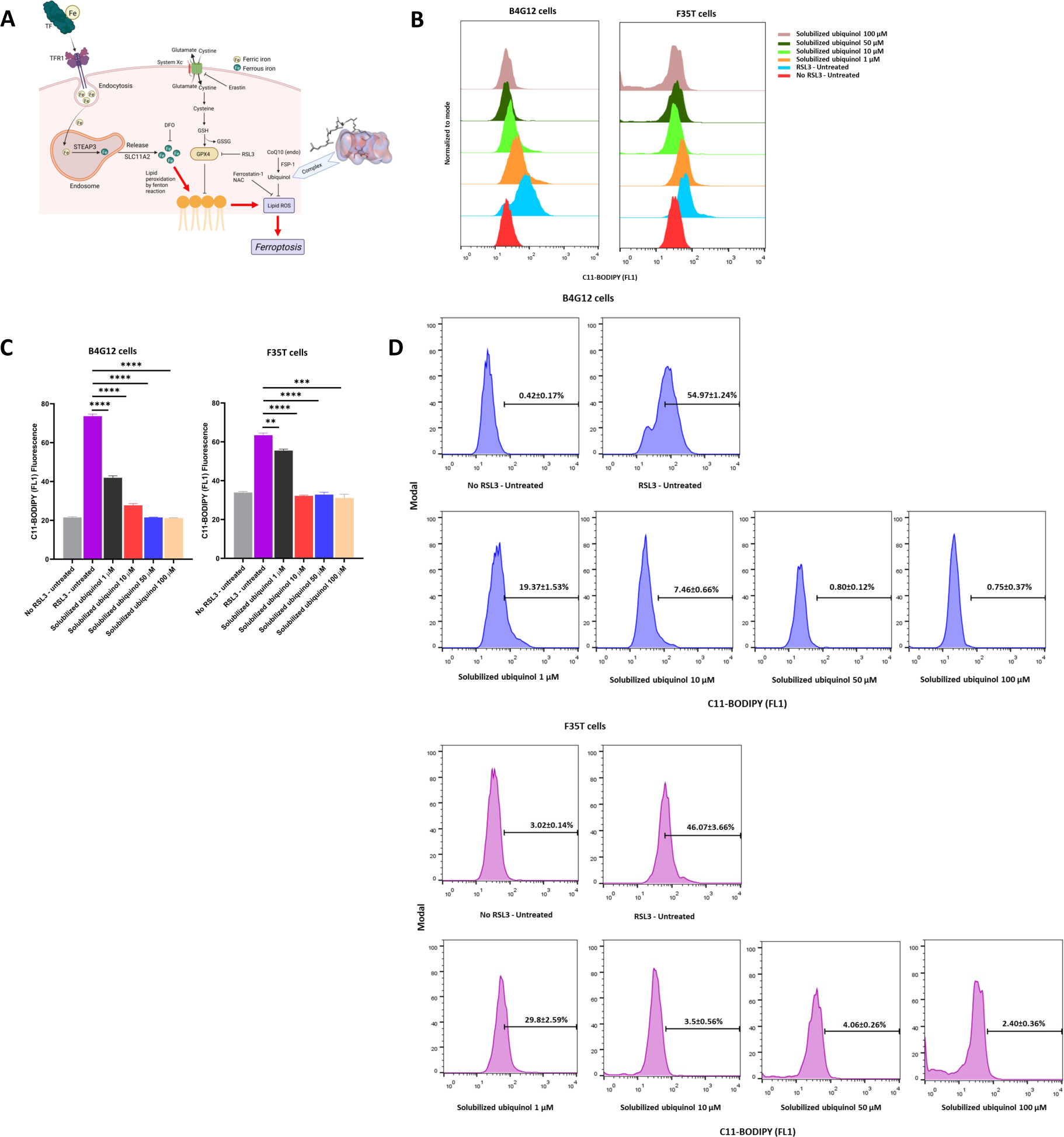
Solubilized ubiquinol gives protection against lipid peroxidation. **(A)** Schematic showing RSL3 induced ferroptosis and role of solubilized ubiquinol to prevent lipid peroxidation and ferroptosis. **(B)** Solubilized ubiquinol prevents lipid peroxidation in HCEC-B4G12 and F35T cells induced by RSL3 in dose dependent manner detected by the peak shift of C11-BODPY fluorescent probe in flow cytometer analysis. **(C)** C11-BODIPY fluorescence signals detected by flow cytometer following treatments with solubilized ubiquinol at different concentrations of 1, 10, 50 and 100 μM. Values are mean±SEM; n = 3; ****p <0.0001, Student’s t-test against RSL3-untreated group. **(D)** Representation of cell population undergoing RSL3 induced lipid peroxidation following solubilized ubiquinol treatment at the indicated concentrations in HCEC-B4G12 and F35T cells. Solubilized ubiquinol decreased the number of cells undergoing lipid peroxidation in a concentration dependent manner. Values are mean±SEM; n = 3.

Interestingly, solubilized ubiquinol treatment provided significant but moderate inhibition of lipid peroxidation at 1 µM in F35T cells with much greater peroxidation reduction in control cells, yet both cell types showed a robust rescue in cell viability (p < 0.0001; **Fig. 6A-D**). This suggests that F35T cells have the ability to survive with higher levels of lipid peroxidation than control cells and that a smaller reduction in lipid peroxidation levels can have a major impact in cell viability. Both cell lines returned to baseline lipid peroxidation levels following 5-10 µM of solubilized ubiquinol treatment (**Fig. 6B-D**). Higher doses of solubilized ubiquinol did not further lower lipid peroxidation levels below baseline but did provide a small increase in cell viability above 100%, suggesting an improvement in cellular health and possibly increased proliferative capacity above basal levels in both cell lines (**Fig. 6C**).

Of note, ubiquinol is an endogenous antioxidant that is converted from coenzyme Q10 via FSP1, a GPX4 glutathione-independent inhibitor of lipid peroxidation and ferroptosis (**Fig. 6A**) (53, 74). Thus, we looked at FSP1 protein expression in our samples and found increased protein expression in both FECD surgical samples (p < 0.05; **Fig. 1B**) and F35T cells (p < 0.001; **Fig. 3M)** compared to controls, indicating endogenous activation of the ubiquinol pathway to help protect against cellular damage and death in FECD.

## DISCUSSION

In this study, we examined ungenotyped FECD patient surgical samples and utilized a variety of immortalized and primary cell culture models from FECD patients with pathological expansions of trinucleotide repeats in intron 3 of the *TCF4* gene (the most common genotype that causes a late onset of disease) to characterize lipid peroxidation and key iron metabolites in FECD and determine whether ferroptosis plays a role in its pathogenesis (49-51). Additionally, we compared changes in iron-lipid interactions that were attributable to genetic background and UVA irradiation against healthy controls, and evaluated the effects of ferroptosis suppressor molecules. We found evidence of elevated lipid peroxide and cytosolic Fe^2+^ levels in FECD samples and models, consistent with increased susceptibility to ferroptosis in FECD attributable to genetic background. We identified the involvement of Fe^2+^, a key mediator of reactive oxidation reactions, in gene-dependent susceptibility to UVA exposure and detected increases in both cytosolic and mitochondrial Fe^2+^ levels after irradiation. TFR1, which has been proposed as a specific ferroptosis marker because it is responsible for the influx of iron inside the cell (36), was found to be elevated in all samples and models and may be a useful biomarker of ferroptosis in this disease. Interestingly, of the inhibitory ferroptosis molecules evaluated (N-acetyl-L-cysteine as a GSH precursor, deferoxamine as a chelator, and ubiquinol as an antioxidant), only solubilized ubiquinol was found to prevent cell death after experimentally induced ferroptosis in this series. Taken altogether, this study presents the first lines of evidence that iron-mediated lipid peroxidation is linked cell death in FECD, and establishes a basis for future mechanistic investigations of ferroptosis prevention in FECD.

While the genetic background of FECD is complex and age of onset varies based on genotype, the clinically recognized hallmarks of this disease are highly conserved. CTG trinucleotide repeat expansions in intron 3 of the *TCF4* gene are found in more than 70% of patients with FECD; patients with this genotypic marker tend to develop “late-onset” disease and tend to be female (3- to 4:1 female to male ratio). Although the exact mechanism of disease remains incompletely understood, it is possible that expressed non-coding regions of mRNA are toxic to CECs. The remaining portion of patients have a mutation in one of at least 10 known genes with a disease-causing mutation that affect a variety of CEC functions, from abnormal collagen production to altered ion channel pump function. Remarkably, all forms of this disease are characterized by an increased susceptibility to oxidative stress, regardless of genotype, age of onset, or gender. Our analysis of surgical tissue explants collected at the time of endothelial keratoplasty for patients with FECD was performed on pooled samples from ungenotyped patients. This broad approach bears direct correlation with the FECD clinical phenotypes common to end-stage disease (loss of CECs, confluent guttae on the inner cornea, corneal edema and vision loss). It is noteworthy that we found evidence of lipid peroxidation, cytosolic Fe^2+^ accumulation, and GPX4 depletion occurring in tandem in surgical samples. Our findings, which are contextualized by the well-documented background of ROS accumulation, support the general presence of increased susceptibility to ferroptosis in patients with FECD. Although we gathered additional immortalized and primary cell culture evidence that the *TCF4* trinucleotide repeat expansion increases ferroptosis susceptibility, further studies are needed to determine if these findings apply to other FECD genotypes equally.

Previously, we showed that ferroptosis can be induced experimentally in healthy immortalized human CECs using erastin, a direct inhibitor of system *x_c_^-^* and the GSH synthetic pathway. In this study, we found evidence that FECD is associated with constitutive increases in cytosolic Fe^2+^ and lipid peroxidation as well as decreased GPX4, specifically in primary and immortalized *in vitro* patient derived FECD cell culture models harboring CTG repeat expansions in *TCF4*. Multiple lines of evidence support the theoretical basis for ferroptosis to be a pathological component of cell death in FECD (4, 12, 26, 75). The *TCF4* genotype models utilized in this study reproduce the ferroptosis pathway signatures found in both diseased tissue and direct experimental induction in healthy cells. Our findings are a strong indication that intronic repeat expansions likely render the FECD endothelium more susceptible to ferroptosis, and underscore the utility of these models – both primary and immortalized FECD cell lines, which returned similar data – in further studies of ferroptosis in FECD. They also suggest possible directions for investigating ferroptosis in other trinucleotide repeat expansions diseases, such as Huntington’s.

In this study elevated levels of Fe^2+^, a reactive redox substrate under strict control and regulation, were detected in the cytosol of CECs from FECD patient samples and cell culture models. While the notion that iron overload plays a part in the pathobiology of Fuchs dystrophy is novel, it is well known that dysregulation of iron metabolism plays a role in several diseases. Typically, iron enters the cell from surrounding fluid after complexation with transferrin, a secretory product that binds Fe^3+^ (**Fig. 7**). The transferrin-Fe^3+^ complex enters via TFR1-mediated endocytosis, becomes acidified and reduced to Fe^2+^ in endosomes by STEAP3 metalloreductase, and gets released into the cytosol where it is converted to Fe^3+^ and incorporated into ferritin, a key iron storage protein and major regulator of intracellular iron (76, 77). Cells normally maintain low free cytosolic Fe^2+^ levels in a steady-state equilibrium with ferritin-bound iron because free Fe^2+^ is highly reactive and potentially toxic. Fe^2+^ can generate excessive ROS via Fenton reactions and increase the activity of lipoxygenases that are responsible for lipid peroxidation (71, 76). Thus, our finding of cytosolic Fe^2+^ accumulation in patient and cell culture models is noteworthy because it implicates ferroptosis as a mechanism of oxidative cell death in FECD. In ferroptosis, the sequence of Fe^2+^-mediated plasma membrane lipid peroxidation and subsequent cell death results in affected cells adopting a characteristic morphological appearance that features redistribution of lipids at the PM, occurring as lipid blebbing and bubble formation and leading toward cellular ballooning prior to death. Although we were unable to visualize ferroptosis cell morphology sequences in tissue explants owing to their terminal degeneration at the end stage of disease, we were able to visualize plasma membrane lipid degeneration and migration of pooled lipids toward the nuclear envelope in cultured FECD cells, confirming ferroptosis cell death sequences of affected cells.

**Figure 7:**
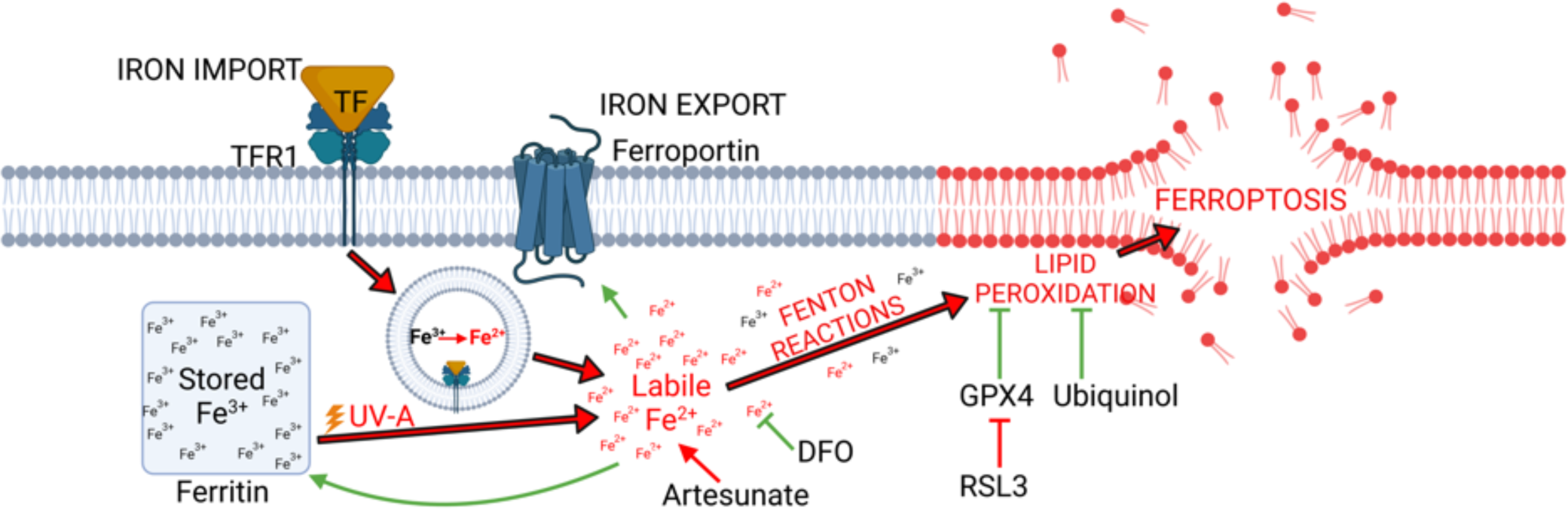
Summary of molecular mechanism of ferroptosis in FECD. Iron enters the cell in ferric form via TFR1-mediated endocytosis. Ferritin stores the excess iron in ferric form, which is nontoxic. The ferric form of iron gets converted to the ferrous form in endosomes. When labile, ferrous iron gets released into the cytosol, it causes lipid peroxidation via Fenton chemistry. UVA irradiation can cause iron release from ferritin which increases the labile iron pool in the cytosol, as well as increases iron-mediated lipid peroxidation, a process known as ferroptosis. GPX4 is the key regulator of ferroptosis, preventing occurrence through scavenging lipid peroxides and reactive oxygen species (ROS). In this study, RSL3 was used to block GPX4 to induce ferroptosis. Ubiquinol, the reduced and active form of coenzyme Q10, is a potent ferroptosis inhibitor that works by scavenging ROS and modulating iron metabolism. Ubiquinol is an essential participant in the FSP1-CoQ10-NAD(P)H pathway, an independent system working in parallel with GPX4 and glutathione to suppress lipid peroxidation and ferroptosis by supporting FSP1 function. Other molecules like DFO and artesunate can prevent ferroptosis by quenching labile toxic ferrous iron, however, are not solely as effective as ubiquinol in preventing ferroptosis.

We detected several points of disruption in iron regulation attributable to FECD genotype, including increased iron uptake signaling and storage disequilibrium. First, cultured FECD CECs with *TCF4* repeat expansions show increased levels of ferroportin, the only membrane bound protein responsible for iron efflux (78). *FPN1* expression has been reported to be regulated by Hif1α, Hif2α and the Hif response element (HRE) under anemic conditions, which would cause increased expression to drive more iron into the blood from cells. More relevant to FECD, *FPN1* expression is also regulated by NRF2 signaling in response to oxidative stress (79), where an increase in ferroportin expression indicates the cellular attempt to remove excess iron from the cytosol as we observed in these experiments. Of note, *FPN1* transcription is also regulated by NRF2 in response to inflammation and in FECD there is evidence of inflammation (80). In addition, increased amounts of transferrin, the secreted iron carrier protein necessary for uptake by cells, were detected in aqueous samples from patients with FECD at the time of transplant surgery. Increased transcription and expression of TFR1, the membrane bound receptor that endocytoses ferritin-Fe^3+^ complexes, was also detected in surgical tissue samples and all cell culture models. As well, decreased levels of cytosolic ferritin were found in cell culture models. The cell behavior observed with respect to iron uptake and storage is consistent with iron scavenging and decreased storage capacity, and these findings may direct future investigations into iron storage regulation and ferritin recycling via ferritinophagy. Interestingly, we also found a corresponding decrease in levels of mitochondrial ferritin, which like its cytosolic counterpart regulates mitochondrial iron and indicates the potential for Fe^2+^ overload in disease-affected CEC mitochondria. Mitochondrial impairment in FECD has been described extensively, with findings that include increased membrane depolarization, decreased ATP synthesis, decreased mtDNA, and increased mitophagy. In this study, we measured mitochondrial iron levels but did not measure mitochondrial membrane lipid peroxidation or other potential indicators of ferroptosis in FECD mitochondria. However, mitochondria may be particularly resistant to ferroptosis due to the presence of a third independent antioxidant defense enzyme, dihydroorotate dehydrogenase (DHODH), that is located on the outer face of the mitochondrial inner membrane and, like FSP1, directly supports the biosynthetic recycling of ubiquinol (81).

We found further evidence of the centrality of Fe^2+^ and importance of lipid peroxidation in FECD disease progression through UVA irradiation and ferroptosis suppression experiments. We first found that increased sensitivity to UV-induced oxidative damage in *TCF4* FECD cell culture models was mediated by Fe^2+^ accumulation. After treating cultured cells with physiologic levels of UVA, we detected increased cytosolic levels of Fe^2+^ in FECD cells compared to controls treated with the same exposures; this response was dose dependent. UVA is a well-known risk factor for FECD disease progression, yet the mechanism by which UV causes oxidative damage in FECD has not been characterized previously. UVA is known to increase cytosolic Fe^2+^ by causing release from ferritin (44, 45, 82). In skin, UVA causes lipid peroxidation in cell membranes via pathways involving Fe (71, 83, 84); ferritin degradation within hours that results in immediate Fe^2+^ increases (82); and compensatory increases in ferritin days after exposure (82, 85). In our study, UVA exposure caused cell death that was nearly identical to RSL3 induced ferroptosis, evidenced by the occurrence of early sharp increase of lipid droplets, dramatic decrease of lipid droplets at the onset of cell death, and presence of lipid droplets at the perinuclear space at the time of death. Our findings support current understandings of the role of ultraviolet as an accelerant of cell damage in FECD by ferroptosis, and demonstrate the involvement of Fe^2+^ in the mechanism of disease progression. We next found that iron chelation with DFO conferred some protection against ferroptosis induced by the direct inhibition of GPX4 using RSL3. When RSL3 inhibits GPX4, the labile iron pool causes lipid peroxidation and increases the accumulation of lipid ROS, resulting ferroptosis mediated cell death (58, 67). Therefore, a larger quantity of labile will increase the risk of ferroptosis when GPX4 is blocked by RSL3. F35T cells had significantly more cell death in comparison to HCEC-B4G12 cells at the same dose of RSL3 challenge and pre-treatment with DFO. DFO was less effective in preventing ferroptosis in F35T cells when compared with HCEC-B4G12 cells due to a comparatively higher labile iron pool. This experiment simultaneously confirmed the presence of increased basal intracellular iron concentrations and susceptibility to cell death in immortalized FECD cells, and provided confirmatory evidence that iron is involved in FECD-associated cell death.

We also found that ferroptosis suppression is possible in FECD cell cultures by co-treatment with ubiquinol. Ubiquinol, the reduced and active form of coenzyme Q10, is an essential participant in the FSP1-CoQ10-NAD(P)H pathway, an independent system working in parallel with GPX4 and glutathione to suppress lipid peroxidation and ferroptosis by supporting FSP1 function (36, 53). This experiment demonstrated the ability to prevent cell death after direct induction of ferroptosis using RSL3 in immortalized FECD cells using a previously validated formulation of ubiquinol complexed with γ-cyclodextrin that increases ubiquinol availability to CECs by increasing drug bioavailability in the aqueous phase. Interestingly, of the inhibitory ferroptosis molecules tested (NAC as a GSH precursor, DFO as a chelator, and ubiquinol as an antioxidant), solubilized ubiquinol clearly outperformed others in preventing cell death at the tested concentrations after experimentally induced ferroptosis. This resonates with our finding of increased FSP1 protein expression indicating endogenous activation of the ubiquinol pathway to help protect against cellular damage and death, and provides further support for ubiquinol targeted therapy in to suppress ferroptosis and cell death in FECD. By demonstrating that specific anti-ferroptosis agents can mitigate the iron-mediated lipid peroxidation present in cells with this common FECD genetic background, we simultaneously confirm the importance of ferroptosis as a mechanism by which characteristic phenotypic damage occurs in CECs and establish the basis for investigating therapeutics to prevent ferroptosis-mediated damage and disease progression in FECD.

Limitations of this study include the use of transformed and expanded primary FECD cell lines. Expanded cell lines propagated from FECD patient samples no longer truly convey the extent of disease *in vitro.* This is due to the growth medium and natural selection for cells that grow most efficiently to confluence, which are most likely the healthiest cells from the donor population. Also, transformed cells are driven to propagate beyond their natural ability. We therefore did not anticipate that all measurements made in this investigation from *in vitro* samples to mirror those from *in vivo* samples. However, we used the cell lines successfully to better understand how the *TCF4* repeat expansion genotype specifically affects ferroptosis susceptibility, and demonstrated that these cell culture models are useful tools for the study of ferroptosis in FECD.

## CONCLUSION

Overall, this investigation presents the first lines of evidence that ferroptosis is a mechanism by which oxidative damage and CEC death occur in Fuchs endothelial corneal dystrophy. We present evidence that expanded trinucleotide repeats in intron 3 of *TCF4* comprise a genetic background that results in increased cytosolic Fe^2+^, iron-mediated lipid peroxidation, and increased susceptibility to ferroptosis. We also present evidence that UVA exposure in cells with expanded repeats in *TCF4* increases the risk of cell death via an iron-mediated mechanism, and highlight the importance of cytosolic Fe^2+^ accumulations as a plausible molecular mechanism for FECD disease progression. Experimental evidence demonstrating that both iron chelation and anti-ferroptosis antioxidant treatments prevent cell death in FECD cell cultures demonstrate the importance of these mechanisms. Although the evidence is strongest in this series for increased ferroptosis susceptibility in this *TCF4* genotype, our findings apply to other FECD genotypes and further studies are needed to explore this possibility.

## MATERIALS AND METHODS

### Consent and tissue collection

All investigations at the University of Iowa and Mayo Clinic were carried out following the guidelines of the Declaration of Helsinki. All tissues were obtained with informed consent by patients or the donor’s family or next of kin. Approval was not required for the deidentified donor corneal tissues in this investigation according to the Institutional Review Board (IRB) at the University of Iowa. For FECD samples, human corneal endothelial tissue was collected at the University of Iowa and Mayo Clinic at the time of endothelial keratoplasty from patients with advanced FECD that were enrolled in the Proteomic Analysis of Corneal Health Study (IRB 201603746) or Mayo Clinic Hereditary Eye Disease Study (IRB 06-007210), respectively. For control samples, human corneal endothelial tissue was obtained from human donor eyes provided by the Iowa Lions Eye Bank (ILEB, Coralville, IA) and Lions Gift of Sight Eye Bank (St. Paul, MN).

### Materials

All required chemicals were procured from commercial sources and utilized without further purification process following the manufacturer’s guidelines. Ubiquinol was procured from Sigma Aldrich (United States Pharmacopeia [USP] reference standard, USA). γ-cyclodextrin was purchased from CI America (Portland, OR). BODIPY™ 581/591 C11 lipid fluorescent probe, Dihydroethidium (Hydroethidine), and SYTOX® Green nucleic acid stain dye were procured from ThermoFisher Scientific (Waltham, MA). Cytosolic ferrous iron (Fe^2+^) detection dye FerroOrange and mitochondrial ferrous iron (Fe^2+^) detection dye Mito-FerroGreen were purchased from Dojindo EU GmbH (Munich, Germany). LipidSpot™ 610 Lipid Droplet Stain was procured from Biotium, Inc., USA. All other solvents and reagents were analytical and cell culture grade.

### Ferroptosis RNASeq data analysis

RNA-Seq datasets of corneal endothelial samples from 47 patients with FECD and 21 donor controls were obtained from publicly available datasets (SRA Accession Numbers: PRJNA445238, PRJNA524323, PRJNA597343). Pre-symptomatic controls with *TCF4* trinucleotide repeats were excluded from the analysis (31, 32, 86). Reads were aligned and gene counts were made using STAR (87), data quality was assessed using FastQC, gene expression was normalized, batch corrected, and determined using EdgeR (88). Genes were excluded if there were less than 3 counts in 20 or more samples. Significant differences (FDR<0.05) were calculated using EdgeR. Gene Set Enrichment Analysis was conducted on known ferroptosis gene signatures (57). For the FerrDB gene signature, all genes that were ferroptosis drivers, suppressors or markers were included, which comprised of 211 expressed genes. Heatmap and hierarchical clustering was conducted using the ComplexHeatMap, cluster, and dendextend packages in R.

### Human Donor and Surgical Tissue Samples

Human donor corneas were procured within 24 hours of donor death and preserved in Optisol-GS storage media (Bausch & Lomb, Irvine, CA) at 4°C. Experiments were conducted within 14 days of preservation. All donors were 50 to 75 years old and each cornea was inspected and evaluated following standard protocols and procedures of the Eye Bank Association of America (EBAA) and ILEB. Endothelial cell-Descemet membrane (EDM) tissues were prepared by mounting donor corneas onto a 9.5 mm vacuum trephine (Barron Precision Instruments, LLC, MI, USA) and scoring the endothelium and Descemet membrane into the stroma. The EDM complex was visualized with 0.06% trypan blue ophthalmic solution (VisionBlue, DORC International, Netherlands) and the tissue was carefully peeled away from the stroma and immediately stored at -80°C until further processing. Human corneal endothelial tissues were collected at the time of endothelial keratoplasty from patients with advanced FECD using standard surgical techniques. Immediately after removal from the eye, approximately 2/3^rd^ of the excised tissue was immediately placed into a cryopreservation vial on dry ice and stored at -80°C until further processing. The remaining 1/3^rd^ of the excised tissue was sent in formalin for histopathological analysis to confirm the diagnosis of FECD.

### Human corneal endothelial cell (HCEC-B4G12) and F35T cell culture

Healthy immortalized human corneal endothelial cells (HCEC-B4G12) were procured from Leibniz Institute DSMZ-German Collection of Microorganisms and Cell Culture GmbH, Germany, and FECD immortalized human corneal endothelial cells (F35T) were a generous gift of Dr. Albert Jun (Johns Hopkins University, Baltimore, MD). F35T cells were derived from a FECD patient expressing the *TCF4* transcript with approximately 4500 CUG repeats in intron 3 (**Supplementary figure 4**). Both cell lines were cultured in Opti-MEM® I Reduced-Serum Media (ThermoFisher) supplemented with 5 ng/mL of human epidermal growth factor (hEGF, ThermoFisher), 20 ng/mL of nerve growth factor (NGF, Fisher Scientific), 200 mg/L of calcium chloride (Sigma-Aldrich), 50 µg/mL of gentamicin (ThermoFisher), 1 mL of Normocin™ (50 mg/mL, Invivogen), 0.08% chondroitin sulfate (Sigma-Aldrich) and 8% fetal bovine serum (HyClone Characterized, US origin). Growth media was optimized for F35T cells and used similarly for B4G12 cells to exclude media-based technical variability. Media was filtered with 0.22 µM PTFE filters prior to use. Cells were incubated at 37°C with a continuous supply of 5% CO_2_ and passaged at the confluence. Plastic surfaces of cell culture dishes were coated with commercial FNC Coating Mix (Athena Environmental Sciences, Inc., USA) to facilitate the adherence of the endothelial cells.

### Real time PCR analysis

Human FECD patient tissues samples (n = 8, multiple tissues pooled together in each group) and donor cornea EDM complexes (n = 4, multiple tissues pooled together in each group) were pelleted, and RNA was extracted and purified using RNeasy kit (Qiagen) according to manufacturer’s instructions. Similarly, HCEC-B4G12 (n = 9) and F35T (n = 6) cells were pelleted, and RNA was extracted. 18 ng total RNA was reverse transcribed using High-Capacity cDNA Reverse Transcription kit (Applied Biosystems). qPCR was performed on the CFX Connect thermal cycler (Bio-Rad Laboratories, Inc) with 10 sec melting, 30 sec annealing/extension for 40 cycles. Melt curve analysis was performed at the end of each qPCR run to verify single product formation. ΔΔCt values were calculated between cell types normalized to *18S* and statistical analysis was performed using Student’s t-test. Primers used for qPCR are mentioned in **supplementary table 8**.

### Western blot analysis

Human FECD patient tissues samples (n = 24 pooled into groups of 3) and donor cornea EDM complexes (n = 12 pooled into groups of 3) were lysed in RIPA buffer with protease inhibitor in pools of 3 tissues for 45 minutes on ice. Similarly, HCEC-B4G12 (n = 5) and F35T (n = 5) cells were pelleted, and protein lysates were extracted. 0.6 µg of total protein was loaded per capillary (DM-TP01, Protein Simple) and lysates were probed with antibodies directed at 4-HNE (STA-035, Cell Biolabs), GPX4 (MAB5457-SP, R&D Systems), NRF2 (PA5-14144, Invitrogen), FSP-1 (20886-1-AP, Proteintech), Ferroportin/SLC40A1 (PA5-G4232, Invitrogen) and TFR1 (MABS1982, Millipore) proteins. Protein expression was normalized to total protein (DM-TP01, Protein Simple) and compared between tissue and cell types.

### Immunohistochemistry for TFR1 protein detection in FECD surgical and control donor corneal tissues

Human surgical explant corneal endothelial tissue was fixed with 4% paraformaldehyde buffered solution, pH 7.4, for 10 minutes at room temperature within 2-4 hours after surgery. Donor corneal endothelial tissue representing a healthy (non-FECD) control sample was fixed following the same protocol within 2 weeks of tissue procurement. Samples were washed three times with 20 mM PBS, pH 7.4, then incubated with 0.1% Triton X-100 in for 30 minutes for permeabilization. Blocking was performed for 1 hour at room temperature in 20 mM PBS, pH 7.4, containing 2% BSA, 5% normal goat serum and 0.1% Triton X-100. Incubation with primary mouse monoclonal anti-TFR1 antibody (clone 3B82A1, catalog# MABS1982, lot# 3519825, EMD Millipore, Burlington, MA, USA) diluted 1:250 with 0.2% BSA, 1% normal goat serum and 0.1% Triton X-100 in 20 mM PBS, pH 7.4 was done for 16 hours at 4°C. Samples were then washed four times with 0.1% Triton X-100 in 20 mM PBS, pH 7.4, and solution containing secondary AlexaFluor568 goat anti-mouse antibody (1:1,000, catalog# A11004, lot# 2447869, Invitrogen, Thermo Fisher Scientific, Waltham, MA, USA) and 300 nM DAPI (Invitrogen, Thermo Fisher Scientific, Waltham, MA, USA) in 0.1% Triton X-100 was applied for two hours at room temperature. Tissue was rinsed with 20 mM PBS, pH 7.4, and distilled water and mounted under Aquamount mounting medium (Thermo Fisher Scientific, Waltham, MA, USA). Images were collected by sequential confocal laser scanning microscopy with a Leica SP8 STED microscope (Leica Microsystems, Mannheim, Germany).

### Cytosolic iron (Fe^2+^) detection in FECD surgical and control donor corneal tissues

FECD corneal tissues were collected during endothelial keratoplasty surgery at the University of Iowa. Immediately after removal from the eye, approximately 2/3^rd^ of the excised tissue was placed in Optisol-GS® and stored at 4°C on wet ice until further processing. Healthy donor corneas stored in Optisol-GS® at 4°C were used to prepare EDM complexes as described above. Human FECD patient tissues samples (n = 7) and donor cornea EDM complexes (n = 11, cut into half and measured as two technical replicates) were digested with 0.2% collagenase type II and 0.05% hyaluronidase in reduced serum OptiMEM-I (Gibco-BRL, Grand Island, NY) media supplemented with 50 µg/mL gentamycin for 3h at 37°C with frequent agitation on a tube rotator (model: 05-450-127, Fisher Scientific). The digestion was completed with 1X 0.5% trypsin-EDTA. Following digestion, cells were filtered with a 100 µM cell strainer to get primary cell suspension. Cells were washed once with Live Imaging Solution, centrifuged, and resuspended in 100 µL Live Imaging Solution. Cells were transferred to a 24-well plate and 300 μL of 1 μmol/L FerroOrange staining solution was added. Cells were incubated for 30 min (37°C, 5% CO_2_) and transferred to flow cytometer tubes. Fluorescence was measured using a flow cytometer (BD FACSCalibur™) and results were analyzed using FlowJo (BD Biosciences, USA).

### Multidimensional protein identification technology (MudPIT) mass spectrometry

Aqueous humor samples from patients with FECD (n = 4) and patients without FECD (n = 4) were collected from patients during surgery. The filter-assisted sample preparation (FASP) method was used for preparing samples for digestion (89). It was solubilized in a mix containing ionic detergent 5% sodium deoxycholate (SDC), buffer 100 mM triethylammonium bicarbonate (TEAB) at pH 8.0, and 3 mM dithiothreitol (DTT). Samples were then sonicated, spun down, and finally the supernatant was transferred to a 30 kD MWCO filter (Millipore, MA, USA) and centrifuged for 30 min at 13,000 g. The filtrate was discarded, and the remaining sample was buffer exchanged with 1% SDC and 100 mM TEAB at pH 8.0. Following buffer exchange, the sample was alkylated with 15 mM iodoacetamide and then digested overnight with trypsin at an enzyme to substrate ratio of 1:100 in a Thermo-Mixer at 1000 RPM at 37°C. Peptides were collected by centrifugation and reversed-phase stop-and-go extraction (STAGE) tips were used for desalting approximately 20 µg of digested peptides (90). Elution solvent was a mixture of 80% acetonitrile and 5% ammonium hydroxide.

Desalted peptides were lyophilized in a SpeedVac (Thermo Fisher Scientific, MA, USA) for 1 h. Peptide samples were then analyzed by ultra-performance liquid chromatography coupled with tandem mass spectrometry (UPLC-MS/MS). The UPLC system was an Easy-nLC 1000 UHPLC system (Thermo Fisher Scientific) coupled with a quadrupole-Orbitrap mass spectrometer (Q-Exactive; Thermo Fisher Scientific). The column was 2 µM Thermo Easy Spray PepMap C18 column (Thermo Fisher Scientific) with 500 mm × 75 µM i.d. Two mobile phases were used, phase A was composed of 97.5% MilliQ water, 2% acetonitrile, and 0.5% formic acid and phase B was composed of 99.5% acetonitrile, and 0.5% formic acid. The elution events were 0-210 min, 0-25% B in A and 210-240 min, 25-80% B in A. Nano-electrospray ionization (Thermo Easy Spray source; Thermo Fisher Scientific) was used at 50°C with an electrospray voltage of 2.2 kV. Tandem mass spectra were acquired from the top 20 ions in the full scan in the range between 400 – 1200 m/z while dynamic exclusion was set to 15 seconds and singly-charged ions were excluded from the analysis. Isolation width was set to 1.6 Dalton and full MS and MS/MS resolution were set to 70,000 and 17,500, respectively. The normalized collision energy was 25 eV and automatic gain control was set to 2e5. Max fill MS and max fill MS/MS were set to 20 and 60 milliseconds, respectively, and the underfill ratio was set to 0.1%.

For identifying peptides, msconvert was used to convert RAW data files to mzML format (91) and then Peak Picker HiRes tool from the OpenMS framework was used to generate MGF files from mzML format (92). Peptide identification searches required precursor mass tolerance of 10 parts per million, fragment mass tolerance of 0.02 Da, strict tryptic cleavage, up to 2 missed cleavages, variable modification of methionine oxidation, fixed modification of cysteine alkylation, and protein-level expectation value scores of 0.0001 or lower. Finally, MGF files were searched using up-to-date protein sequence libraries available from X!Tandem (93), UniProtKB, and OMSSA (94). Identified protein intensities were normalized to total peptide hits per sample and scaled to logarithmic base 10. Partek Genomics Suite version 7.21.1119. (MO, USA) was used to determine statistically significant proteins (analysis of variance [ANOVA], p < 0.05). Pathway significance was ascertained by the number of proteins in the dataset in common with known proteins in a single pathway, as determined by the IPA database (Qiagen).

### Basal level of ROS quantification

HCEC-B4G12 and F35T cells at 250,000 cells/well in 24-well plate were stained with 10 µM of dihydroethidium (DHE) in 1 mL of Live Imaging Solution and incubated for 30 min (37°C, 5% CO_2_). After incubation, cells were transferred to flow cytometer tubes. Fluorescence was measured using a flow cytometer (BD FACSCalibur) and data was analyzed using FlowJo.

### Confocal microscopy

Confocal microscopy was conducted with multiple fluorescent probes, including the Dihydroethidium (Hydroethidine; DHE) for fluorescent probe for imaging ROS, C11-Bodipy 581/591 for lipid peroxidation, FerroOrange for cytosolic labile iron, Mito-FerroGreen for mitochondrial iron, and LipidSpot™ 610 for lipid droplets. HCEC-B4G12 and F35T cells were seeded at 100,000 to 200,000 cells/chamber in 4-chambered coverglass slides (Thermo Scientific Nunc Lab-Tek) after coating the glass surface with FNC Coating Mix and incubating at 37°C and 5% CO_2_ 18 to 72 h. After incubation, media was discarded, and cells were washed with Live Imaging Solution if necessary. For iron detection, washing at least twice was important to remove extracellular iron from the media. For lipid peroxidation imaging, cells were stained with 0.9 mL of 5 µM C11-Bodipy 581/591 fluorescent probe in Live Imaging Solution for 20 min and stain solution was discarded before adding 0.9 mL of fresh Live Imaging Solution. For cytosolic and mitochondrial iron imaging, after washing twice, 500 µL of 1 μmol/L FerroOrange and 5 µmol/L Mito-FerroGreen staining solutions were added to the cells and incubated for 30 min, separately. For ROS imaging, cells were stained with 0.9 mL of 10 µM Dihydroethidium fluorescent probe in Live Imaging Solution for 30 min. For lipid droplet imaging, cells were stained with 0.9 mL of 1X LipidSpot™ 610 fluorescent probe in cell culture media for 30 min. All incubations were conducted in a cell culture incubator at 37°C and 5% CO_2_. Immediately after incubation, live cells were imaged using a Leica SP8 confocal microscope with a 63X oil lens equipped with Leica Application Suite X (LAS X) operating software. For each filter, all images were taken at the same gain level and image capture settings for both HCEC-B4G12 and F35T cells. Images were analyzed using Image J.

### Mitochondrial superoxide assay

HCEC-B4G12 (n = 12) and F35T (n = 8) cells (50,000/well and 25,000/well, respectively) were grown in 96 well plates until reaching confluency. Cells were exposed to MitoROS 580 dye (ab219943, Abcam) for 30 min, and mitochondrial superoxide was quantified using Infinite M Plex plate reader (Tecan Group, Ltd) with Ex/Em =540/590 nm. As a positive assay control, additional B4G12 and F35T cells were treated with antimycin-A (AMA) for 30 min prior to and during the MitoROS 580 dye treatment, for a total of 60 min. Cells were fixed with 4% formaldehyde buffered in 0.1 M PBS, pH 7.4 for 30 min, washed with PBS 3 times, and incubated with 300 nM DAPI for 30 min. DAPI-stained nuclei were counted in whole well images captured by Cytation 5 instrument (BioTek Instruments, Inc) using Gen5 Image Plus (version 3.10) software, and resulting cell counts were applied to obtain normalized MitoROS 580 RFU/cell values for each well. Average values were calculated per group and compared using Student’s t-test.

### Cytosolic iron (Fe^2+^) detection by flow cytometry using FerroOrange fluorescent probe

HCEC-B4G12 and F35T cells were cultured in T75 flasks until they reached confluency. Cells were trypsinized and washed twice with 1X DPBS buffer to remove residual trypsin and serum-containing media. The cell suspension was centrifuged at 230 g for 5 min and resuspended in Live Imaging Solution (Thermo Fisher). FerroOrange staining solution of 1 μmol/L was prepared following the manufacturer’s guidelines. In 24 well plates, 150,000 cells in 100 µL Live Imaging Solution were added to each well (n = 3) and 300 µL of 1 μmol/L FerroOrange staining solution was added. For the control group, 300 µL of Live Imaging Solution was added to the cells (n = 3). Then cells were incubated for 30 min (37°C, 5% CO_2_). After incubation, cells were transferred to flow cytometry tubes and fluorescence was measured by flow cytometry (BD FACSCalibur™). Data analysis was carried out using FlowJo software.

### Mitochondrial iron (Fe^2+^) detection by flow cytometry using Mito-FerroGreen fluorescent probe

HCEC-B4G12 and F35T cells were cultured, trypsinized, washed, and resuspended in Live Imaging Solution following the same protocol as cytosolic iron detection. Mito-FerroGreen staining solution of 5 μmol/L was prepared following the manufacturer’s guidelines. For the experimental group, 500 µL of 5 µmol/L Mito-FerroGreen staining solution was added to 150,000 cells in 100 µL of Live Imaging Solution in a 24-well plate (n = 3). For the control group, 500 µL of Live Imaging Solution was added to the cells (n = 3). After 30 min incubation (37°C, 5% CO_2_), fluorescence was measured using a flow cytometer (BD FACSCalibur™) and data was analyzed using FlowJo.

### Cytosolic ferritin ELISA

HCEC-B4G12 and F35T cells were cultured in T75 flasks with 3 biological replicates. At confluence, the protein was extracted with 1X RIPA buffer (Sigma-Aldrich) supplied with EDTA-free Protease Inhibitor Cocktail (cOmplete™, Roche). Protein was stored at -80°C until the ELISA assay was performed. Protein concentration in each biological replicate was measured by Pierce™ BCA Protein Assay Kit (Thermo Scientific™). Three technical replicates of 100 µg protein from each biological replicate diluted with diluent supplied by the manufacturer were added to the antibody-coated wells (Ferritin Human ELISA Kit, Invitrogen). All the steps were carried out following the manufacturer’s protocol. Spectra Max plus 384 Microplate Spectrophotometer was used to measure the absorbance at 490 nm. The standard calibration curve of ferritin was prepared in duplicates.

### Mitochondrial ferritin (MTFT) ELISA

Mitochondrial ferritin in HCEC-B4G12 and F35T cells were quantified using Immunotag™ Mitochondrial Ferritin ELISA kit (G-Biosciences, USA). Protein was extracted following the same protocol used for the cytosolic ferritin ELISA. Three technical replicates of 100 µg protein from each biological replicate (n = 3) were added to antibody-coated wells and all the experimental steps were conducted according to the manufacturer’s guidelines without any modification. The standard calibration curve of FTMT was prepared in duplicate.

### Basal level of lipid peroxidation (C11-Bodipy 581/591) assay in HCEC-B4G12 and F35T cells

HCEC-B4G12 and F35T cells were cultured in T75 flasks until they reached confluency. After detachment with trypsin, 400,000 cells were stained with 2 mL of 5 µM C11-Bodipy 581/591 fluorescent probe in Live Imaging Solution (Thermo Fisher) or left unstained (unstained control group), mixed by pipetting, and incubated in a cell culture incubator (37°C and 5% CO_2_) for 20 min. After staining, the cell suspension was transferred to 15 mL Falcon tubes. All 15 mL tubes were centrifuged at 230 g for 5 min, then stain solution was discarded, and the cells were resuspended in 0.6 mL of Live Imaging Solution. After cells were transferred to flow cytometer tubes, fluorescence was measured using a flow cytometer (BD FACSCalibur™) and results were analyzed using FlowJo.

### Quantification of lipid droplets

HCEC-B4G12 and F35T cells at 250,000 cells/well in 24-well plate were stained with 1X of LipidSpot™ 610 in 1 mL of cell culture media and incubated for 30 min (37°C, 5% CO_2_). Following incubation, cells were transferred to flow cytometer tubes. Fluorescence was measured using a flow cytometer (BD FACSCalibur) and data was analyzed using FlowJo.

### Time-lapse confocal imaging

HCEC-B4G12 and F35T cells were seeded at 100,000-150,000 cells/chamber in a 4-chambered glass bottom coverglass slide (Thermo Scientific Nunc Lab-Tek) after coating with FNC Coating Mix and incubated for 18 h (37°C and 5% CO_2_). Cells were treated with 1 µM of RSL3, 1X of LipidSpot™ 610 and 50 nM of SYTOX™ Green nucleic acid stain dye in 0.9 mL of cell culture media. In another experiment, cells were exposed with UVA at 1.5 J/cm^2^. Following UVA exposure, cells were treated with 1X of LipidSpot™ 610 and 50 nM of SYTOX™ Green nucleic acid stain dye in 0.9 mL of cell culture media. Time-lapse Z-stack imaging was performed for 17 h to 48 h using a LSM 980 confocal microscope (Zeiss) with Airyscan 2 with 63X oil lens. During imaging, cells incubated in the chamber were maintained at 37°C and 5% CO_2_. Image analysis and video production were performed using Imaris 9.9 (Oxford Instruments).

### Deferoxamine (DFO) iron chelation assay

HCEC-B4G12 and F35T cells were seeded in 96-well plates at 5,000 cells/well. After 18 h of incubation at 37°C and a continuous supply of 5% CO_2_, cells were treated with 100 µM of DFO (n = 3). After incubating for 24 h (37°C and 5% CO_2_), cells were washed once with 1X DPBS, and RSL3 at the doses of 1, 2, and 5 µM in DMSO were added to the respective cells while an equal amount of DMSO was added to the control cells, and incubated for 2, 4, 6 and 8 h (37°C and 5% CO_2_). After the RSL3 treatment of 2, 4, 6, and 8 h, at each time point, cells were gently washed once with 1X DPBS and MTS reagent of 20 μL (Cell Titer-96 Aqueous One Solution, Promega, USA) in 80 μL of cell culture media for 3 h (37°C and 5% CO_2_). After the incubation, the absorbance was measured at 490 nm using the Spectra Max plus 384 Microplate Spectrophotometer (Molecular Devices, Sunnyvale, CA) following the manufacturer’s guidelines.

### Preparation of solubilized ubiquinol

Solubilized ubiquinol was prepared by the kneading method where physical complexation was formed between ubiquinol and γ-cyclodextrin (γ-CD) following our previously published protocol (33). Briefly, ubiquinol and γ-CD were mixed at the molar ratio of 1:10 and a hydro-alcoholic solution at 1:1 ratio was added to the mixture to form a semi-liquid paste. Mixing was continued for 1 h and then the paste was vacuum dried to yield the powdered complex.

### Lipid peroxidation inhibition by solubilized ubiquinol in HCEC-B4G12 and F35T cells

The lipid peroxidation assay was conducted using C11-Bodipy 581/591 fluorescent probe following the same seeding, washing, staining and measurement steps described in the basal level of lipid peroxidation assay, except cells were treated with solubilized ubiquinol then challenged with 1 µM of RSL3 or left untreated as controls. 18 h after seeding, media was removed and solubilized ubiquinol at concentrations of 1, 10, 50, and 100 µM diluted in cell culture media were added to the cells (n = 3) and incubated for 24 h (37°C and 5% CO_2_). After incubation, cells were washed twice with 1X DPBS, and 1 µM of RSL3 in DMSO was added except for the untreated and untreated-unstained control groups. Cells were then incubated for 8 h at 37°C and 5% CO_2_, and after the incubation, cells were stained with 2 µL of C11-Bodipy 581/591 fluorescent probe stock prepared in DMSO as described in the basal lipid peroxidation assay.

### Ferroptosis assay in HCEC-B4G12 and F35T cells

Lactate dehydrogenase (LDH) assay was performed using CyQUANT™ LDH Cytotoxicity Assay kit (Invitrogen, USA). LDH release in media was quantified as an endpoint of measuring ferroptosis. HCEC-B4G12 and F35T cells were seeded at 5,000 cells/well in a 96-well plate and incubated for 18 h (37°C and 5% CO_2_). After incubation, media was removed, and solubilized ubiquinol dispersed in media was added at 1, 5, 10, 50, and 100 µM concentrations while only media was added to the control group. In this assay, one set of cells was used for measuring the spontaneous LDH activity, and another set of cells was used for measuring maximum LDH activity. Ferrostatin-1 (Sigma-Aldrich, USA), a known ferroptosis inhibitor, was dissolved in DMSO at 1 µM and used as an anti-ferroptotic positive control. After incubating for 24 h at 37°C and 5% CO_2_, cells were washed twice with 1X DPBS and treated with 1 µM of RSL3 in DMSO whereas control groups were only treated with an equal amount of DMSO in media. After 24 h of incubation, 10 μL of supplied 10X lysis buffer was added to the group designated for measuring the maximum LDH activity and incubated in the cell culture incubator for 45 min. Following 45 min incubation, 50 µL from each well was transferred to a new 96-well flat-bottom well plate. To test the assay performance, 50 μL of 1X LDH positive control was added to three wells which were used as the LDH positive control group. 50 μL of supplied reaction mixture was added and incubated for 30 min in the dark at room temperature (RT). 50 μL of stop solution was added to each well to stop the reaction. Absorbance was measured at 490 nm following the manufacturer’s protocol. Percent cell viability was calculated using the formula mentioned below:

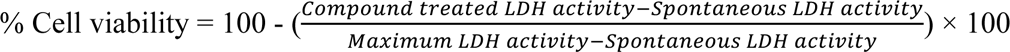

### Ferroptosis assay in F35T cells using MTS assay

F35T cells were seeded at 2,500 and 5,000 cells/well in 96-well plate and incubated for 18 h (37°C and 5% CO_2_). After the incubation, media was removed, and solubilized ubiquinol, N-Acetylcysteine (NAC) and deferoxamine (DFO) dispersed in media were added separately at 1 and 10 µM concentrations while only media was added to the control group. Ferroptosis inhibitor ferrostatin-1 (Sigma-Aldrich, USA) in DMSO was used as a positive control. After adding treatments, cells were incubated for 24 h (37°C and 5% CO_2_). Cells were then washed twice with 1X DPBS to remove any residual treatments and 1 µM of RSL3 in DMSO was added to the cells except for the no RSL3 control group where only DMSO in media was added. After adding RSL3, cells were incubated for an additional 24 h (37°C and 5% CO_2_) and bright-field microscopic images of live cells were taken using the EVOS Cell Imaging System (ThermoFisher, USA). Cells were gently washed once with 1X DPBS and 20 μL of MTS reagent in 80 μL of media was added to the cells and incubated for 3 h (37°C and 5% CO_2_). After the incubation, the absorbance was measured at 490 nm following the manufacturer’s guidelines.

### Cytosolic iron (Fe^2+^) detection in human donor corneas upon UVA irradiation

Human donor corneas stored in Optisol-GS® at 4°C were procured as noted above. This experiment was performed in pairwise fashion, where the right eye was exposed to UVA irradiation (n = 3) and the left eye from the same donor was used as a control. Corneas were washed with sterile Hank’s balanced salt solution (HBSS) and placed in 12-well plate in HBSS with endothelial side facing the UVA light source. Cells were exposed to UVA irradiation at the fluence of 5 J/cm^2^ using a Rayonet Photochemical Reactor (RPR-200, The Southern NE Ultraviolet Co., Brandford, CT). Initially, various UVA fluence levels of 5 to 25 J/cm^2^ with 5 J/cm^2^ increments were screened to find a safe UVA dose for cells by measuring cell viability after UVA irradiation using a MTS assay kit. In our UVA dose screening, we observed that doses above 5 J/cm^2^ are toxic to F35T cells; thus, a dose of 5 J/cm^2^ was chosen as a relatively safe UVA dose for measuring cell viability and sensitivity to UVA irradiation. Control corneas were treated identically except they were not subjected to UVA irradiation. Both UVA irradiated and non-irradiated control corneas were digested following the same procedures noted previously and primary cell suspensions were obtained. Cells were resuspended in 100 µL of Live Imaging Solution and transferred to a 24-well plate and 600 μL of 1 μmol/L FerroOrange staining solution was added to each well. Staining was performed for 30 min (37°C and 5% CO_2_). Fluorescence was measured using a flow cytometer (BD FACSCalibur) and data were analyzed using FloJo.

### Cytosolic and mitochondrial Iron (Fe^2+^) detection following UVA irradiation

HCEC-B4G12 and F35T cells were cultured and prepared following the same protocol as used for cytosolic iron detection. To quantify cytosolic iron, 400,000 cells in 300 µL of Live Imaging Solution were added to each well in a 24 well plate. The UVA irradiation instrument set-up was the same as noted previously for the irradiation of human corneas. Various doses of UVA irradiation of 1, 2, 4, and 8 J/cm^2^ fluences were delivered. UVA irradiated and non-irradiated control cells were stained with 1.2 mL of 1 μmol/L FerroOrange staining solution for 30 min in the incubator (37°C, 5% CO_2_). For the detection of mitochondrial iron, 20,000 cells in 200 µL of Live Imaging Solution were added to each well and irradiated with a UV dose of 1, 2, 4, and 8 J/cm^2^. UV irradiated and non-irradiated control cells were stained with 700 µL of 5 μmol/L Mito-FerroGreen staining solution for 30 min in an incubator (37°C, 5% CO_2_). The same volume of Live Imaging solution was added to the unstained control group, which was used to monitor background fluorescence signals. After incubation, fluorescence was measured using a flow cytometer (BD FACSCalibur) and data was analyzed using FlowJo.

### Statistical analysis

All data were expressed as the mean ± standard error of the mean (SEM). Statistical analysis was performed using the two-tailed Student’s t-test when the experimental group was only compared with the control group. One-way ANOVA followed by Tukey’s post-hoc test was utilized when multiple groups were compared with each other. P-values of less than 0.05 were considered statistically significant. All experiments were carried out with at least 3 biological replicates and in technical triplicate. Statistical analysis was carried out using GraphPad Prism.

## Supporting information

STable 5A

STable 5B

STable 6

## Acknowledgements

The authors cordially thank the Lyle and Sharon Bighley Endowed Chair of Pharmaceutical Sciences, Iowa Lions Eye Bank, the Beulah and Florence Usher Endowed Chair in Cornea/External Disease and Refractive Surgery, the M.D. Wagoner & M.A. Greiner Cornea Excellence Fund, Mr. and Mrs. Robert and Joell Brightfelt, Mr. and Mrs. Lloyd and Betty Schermer, the UIHC Cornea Research Fund for financial support, and the cornea patients, donors, and donor families that made this research possible. MAG, AKS, JMS and SS are supported by the NIH grant R21EY034198.

## List of abbreviations

DFO: Deferoxamine
DHE: Dihydroethidium
EDM: Endothelial cell-Descemet membrane
FASP: Filter assisted sample preparation
FECD: Fuchs endothelial corneal dystrophy
ILEB: Iowa Lions Eye Bank
IRB: Institutional Review Board
LDH: Lactate dehydrogenase
MudPIT: Multidimensional protein identification technology
NAC: N-Acetylcysteine
NGF: Nerve growth factor
ROS: Reactive oxygen species
SEM: Standard error of the mean
TFRA: transferrin receptor 1
USP: United States Pharmacopeia
UVA: Ultraviolet A

## References

1. Ołdak M, Ruszkowska E, Udziela M, Oziębło D, Bińczyk E, Ścieżyńska A, et al. Fuchs Endothelial Corneal Dystrophy: Strong Association with rs613872 Not Paralleled by Changes in Corneal Endothelial TCF4 mRNA Level. BioMed research international. 2015;2015:640234.

2. Uchida T, Sakai O, Imai H, Ueta T. Role of Glutathione Peroxidase 4 in Corneal Endothelial Cells. Curr Eye Res. 2017;42(3):380–5.

3. Lovatt M, Adnan K, Peh GSL, Mehta JS. Regulation of Oxidative Stress in Corneal Endothelial Cells by Prdx6. Antioxidants (Basel). 2018;7(12).

4. Lovatt M, Adnan K, Kocaba V, Dirisamer M, Peh GSL, Mehta JS. Peroxiredoxin-1 regulates lipid peroxidation in corneal endothelial cells. Redox biology. 2020;30:101417.

5. Eye Bank Association of America. 2019 Eye Banking Statistical Report. Washington, DC; 2020.

6. Wagoner MD, Bohrer LR, Aldrich BT, Greiner MA, Mullins RF, Worthington KS, et al. Feeder-free differentiation of cells exhibiting characteristics of corneal endothelium from human induced pluripotent stem cells. Biol Open. 2018;7(5).

7. Joyce NC. Proliferative capacity of corneal endothelial cells. Exp Eye Res. 2012;95(1):16–23.

8. Jalimarada SS, Ogando DG, Bonanno JA. Loss of ion transporters and increased unfolded protein response in Fuchs’ dystrophy. Mol Vis. 2014;20:1668–79.

9. Liu C, Miyajima T, Melangath G, Miyai T, Vasanth S, Deshpande N, et al. Ultraviolet A light induces DNA damage and estrogen-DNA adducts in Fuchs endothelial corneal dystrophy causing females to be more affected. Proc Natl Acad Sci U S A. 2020;117(1):573–83.

10. Liu C, Vojnovic D, Kochevar IE, Jurkunas UV. UV-A Irradiation Activates Nrf2-Regulated Antioxidant Defense and Induces p53/Caspase3-Dependent Apoptosis in Corneal Endothelial Cells. Invest Ophthalmol Vis Sci. 2016;57(4):2319–27.

11. White TL, Deshpande N, Kumar V, Gauthier AG, Jurkunas UV. Cell cycle re-entry and arrest in G2/M phase induces senescence and fibrosis in Fuchs Endothelial Corneal Dystrophy. Free Radic Biol Med. 2021;164:34–43.

12. Jurkunas UV, Bitar MS, Funaki T, Azizi B. Evidence of oxidative stress in the pathogenesis of fuchs endothelial corneal dystrophy. The American journal of pathology. 2010;177(5):2278–89.

13. Zinflou C, Rochette PJ. Ultraviolet A-induced oxidation in cornea: Characterization of the early oxidation-related events. Free Radic Biol Med. 2017;108:118–28.

14. Delic NC, Lyons JG, Di Girolamo N, Halliday GM. Damaging Effects of Ultraviolet Radiation on the Cornea. Photochem Photobiol. 2017;93(4):920–9.

15. Czarny P, Kasprzak E, Wielgorski M, Udziela M, Markiewicz B, Blasiak J, et al. DNA damage and repair in Fuchs endothelial corneal dystrophy. Mol Biol Rep. 2013;40(4):2977–83.

16. Halilovic A, Schmedt T, Benischke AS, Hamill C, Chen Y, Santos JH, et al. Menadione-Induced DNA Damage Leads to Mitochondrial Dysfunction and Fragmentation During Rosette Formation in Fuchs Endothelial Corneal Dystrophy. Antioxidants & redox signaling. 2016;24(18):1072–83.

17. Kumar V, Jurkunas UV. Mitochondrial Dysfunction and Mitophagy in Fuchs Endothelial Corneal Dystrophy. Cells. 2021;10(8).

18. Moysan A, Marquis I, Gaboriau F, Santus R, Dubertret L, Morlière P. Ultraviolet A-induced lipid peroxidation and antioxidant defense systems in cultured human skin fibroblasts. J Invest Dermatol. 1993;100(5):692–8.

19. Vats K, Kruglov O, Mizes A, Samovich SN, Amoscato AA, Tyurin VA, et al. Keratinocyte death by ferroptosis initiates skin inflammation after UVB exposure. Redox Biology. 2021;47:102143.

20. Jurkunas UV, Bitar MS, Funaki T, Azizi B. Evidence of oxidative stress in the pathogenesis of fuchs endothelial corneal dystrophy. Am J Pathol. 2010;177(5):2278–89.

21. Jurkunas UV, Rawe I, Bitar MS, Zhu C, Harris DL, Colby K, et al. Decreased expression of peroxiredoxins in Fuchs’ endothelial dystrophy. Invest Ophthalmol Vis Sci. 2008;49(7):2956–63.

22. Gottsch JD, Bowers AL, Margulies EH, Seitzman GD, Kim SW, Saha S, et al. Serial analysis of gene expression in the corneal endothelium of Fuchs’ dystrophy. Invest Ophthalmol Vis Sci. 2003;44(2):594–9.

23. Buddi R, Lin B, Atilano SR, Zorapapel NC, Kenney MC, Brown DJ. Evidence of oxidative stress in human corneal diseases. J Histochem Cytochem. 2002;50(3):341–51.

24. Lovatt M, Kocaba V, Hui Neo DJ, Soh YQ, Mehta JS. Nrf2: A unifying transcription factor in the pathogenesis of Fuchs’ endothelial corneal dystrophy. Redox Biology. 2020;37:101763.

25. Kuang F, Liu J, Tang D, Kang R. Oxidative Damage and Antioxidant Defense in Ferroptosis. Frontiers in cell and developmental biology. 2020;8:586578.

26. Liu C, Chen Y, Kochevar IE, Jurkunas UV. Decreased DJ-1 leads to impaired Nrf2-regulated antioxidant defense and increased UV-A-induced apoptosis in corneal endothelial cells. Invest Ophthalmol Vis Sci. 2014;55(9):5551–60.

27. Dodson M, Castro-Portuguez R, Zhang DD. NRF2 plays a critical role in mitigating lipid peroxidation and ferroptosis. Redox Biology. 2019;23:101107.

28. Yang Wan S, SriRamaratnam R, Welsch Matthew E, Shimada K, Skouta R, Viswanathan Vasanthi S, et al. Regulation of Ferroptotic Cancer Cell Death by GPX4. Cell. 2014;156(1):317–31.

29. Forcina GC, Dixon SJ. GPX4 at the Crossroads of Lipid Homeostasis and Ferroptosis. Proteomics. 2019;19(18):e1800311.

30. Yang WS, Stockwell BR. Synthetic Lethal Screening Identifies Compounds Activating Iron-Dependent, Nonapoptotic Cell Death in Oncogenic-RAS-Harboring Cancer Cells. Chem Biol. 2008;15(3):234–45.

31. Nikitina AS, Belodedova AV, Malyugin BE, Sharova EI, Kostryukova ES, Larin AK, et al. Dataset on transcriptome profiling of corneal endothelium from patients with Fuchs endothelial corneal dystrophy. Data in brief. 2019;25:104047.

32. Wieben ED, Aleff RA, Tang X, Kalari KR, Maguire LJ, Patel SV, et al. Gene expression in the corneal endothelium of Fuchs endothelial corneal dystrophy patients with and without expansion of a trinucleotide repeat in TCF4. PLoS One. 2018;13(7):e0200005.

33. Naguib YW, Saha S, Skeie JM, Acri T, Ebeid K, Abdel-rahman S, et al. Solubilized ubiquinol for preserving corneal function. Biomaterials. 2021;275:120842.

34. Dixon SJ, Lemberg KM, Lamprecht MR, Skouta R, Zaitsev EM, Gleason CE, et al. Ferroptosis: an iron-dependent form of nonapoptotic cell death. Cell. 2012;149(5):1060–72.

35. Li J, Cao F, Yin H-l, Huang Z-j, Lin Z-t, Mao N, et al. Ferroptosis: past, present and future. Cell Death Dis. 2020;11(2):88.

36. Feng H, Schorpp K, Jin J, Yozwiak CE, Hoffstrom BG, Decker AM, et al. Transferrin Receptor Is a Specific Ferroptosis Marker. Cell Rep. 2020;30(10):3411–23.

37. Bogdan AR, Miyazawa M, Hashimoto K, Tsuji Y. Regulators of Iron Homeostasis: New Players in Metabolism, Cell Death, and Disease. Trends Biochem Sci. 2016;41(3):274–86.

38. Venkataramani V. Iron Homeostasis and Metabolism: Two Sides of a Coin. Adv Exp Med Biol. 2021;1301:25–40.

39. Girotti AW. Lipid hydroperoxide generation, turnover, and effector action in biological systems. J Lipid Res. 1998;39(8):1529–42.

40. Girotti AW, Kriska T. Role of lipid hydroperoxides in photo-oxidative stress signaling. Antioxid Redox Signal. 2004;6(2):301–10.

41. Aust SD, Morehouse LA, Thomas CE. Role of metals in oxygen radical reactions. J Free Radic Biol Med. 1985;1(1):3–25.

42. Clemente SM, Martínez-Costa OH, Monsalve M, Samhan-Arias AK. Targeting Lipid Peroxidation for Cancer Treatment. Molecules. 2020;25(21).

43. Halliwell B, Gutteridge JM. Biologically relevant metal ion-dependent hydroxyl radical generation. An update. FEBS Lett. 1992;307(1):108–12.

44. Aubailly M, Santus R, Salmon S. Ferrous ion release from ferritin by ultraviolet-A radiations. Photochem Photobiol. 1991;54(5):769–73.

45. Wolszczak M, Gajda J. Iron release from ferritin induced by light and ionizing radiation. Research on Chemical Intermediates. 2010;36(5):549–63.

46. Zhu J, Dai P, Liu F, Li Y, Qin Y, Yang Q, et al. Upconverting Nanocarriers Enable Triggered Microtubule Inhibition and Concurrent Ferroptosis Induction for Selective Treatment of Triple-Negative Breast Cancer. Nano Lett. 2020;20(9):6235–45.

47. Smith MJ, Fowler M, Naftalin RJ, Siow RCM. UVA irradiation increases ferrous iron release from human skin fibroblast and endothelial cell ferritin: Consequences for cell senescence and aging. Free Radic Biol Med. 2020;155:49–57.

48. Vile GF, Tyrrell RM. Oxidative stress resulting from ultraviolet A irradiation of human skin fibroblasts leads to a heme oxygenase-dependent increase in ferritin. J Biol Chem. 1993;268(20):14678–81.

49. Afshari NA, Igo RP, Morris NJ, Stambolian D, Sharma S, Pulagam VL, et al. Genome-wide association study identifies three novel loci in Fuchs endothelial corneal dystrophy. Nature Communications. 2017;8(1):14898.

50. Wieben ED, Aleff RA, Rinkoski TA, Baratz KH, Basu S, Patel SV, et al. Comparison of TCF4 repeat expansion length in corneal endothelium and leukocytes of patients with Fuchs endothelial corneal dystrophy. PLoS One. 2021;16(12):e0260837.

51. Fautsch MP, Wieben ED, Baratz KH, Bhattacharyya N, Sadan AN, Hafford-Tear NJ, et al. TCF4-mediated Fuchs endothelial corneal dystrophy: Insights into a common trinucleotide repeat-associated disease. Prog Retin Eye Res. 2021;81:100883.

52. Chen X, Yu C, Kang R, Tang D. Iron Metabolism in Ferroptosis. 2020;8.

53. Doll S, Freitas FP, Shah R, Aldrovandi M, da Silva MC, Ingold I, et al. FSP1 is a glutathione-independent ferroptosis suppressor. Nature. 2019;575(7784):693-8.

54. Hu W, Zhou C, Jing Q, Li Y, Yang J, Yang C, et al. FTH promotes the proliferation and renders the HCC cells specifically resist to ferroptosis by maintaining iron homeostasis. Cancer Cell International. 2021;21(1):709.

55. Ke S, Wang C, Su Z, Lin S, Wu G. Integrated Analysis Reveals Critical Ferroptosis Regulators and FTL Contribute to Cancer Progression in Hepatocellular Carcinoma. 2022;13.

56. Seibt TM, Proneth B, Conrad M. Role of GPX4 in ferroptosis and its pharmacological implication. Free Radical Biology and Medicine. 2019;133:144–52.

57. Zhou N, Bao J. FerrDb: a manually curated resource for regulators and markers of ferroptosis and ferroptosis-disease associations. Database: the journal of biological databases and curation. 2020;2020.

58. Dixon Scott J, Lemberg Kathryn M, Lamprecht Michael R, Skouta R, Zaitsev Eleina M, Gleason Caroline E, et al. Ferroptosis: An Iron-Dependent Form of Nonapoptotic Cell Death. Cell. 2012;149(5):1060–72.

59. Kerins MJ, Ooi A. The Roles of NRF2 in Modulating Cellular Iron Homeostasis. Antioxid Redox Signal. 2018;29(17):1756–73.

60. Fortenbach CR, Skeie JM, Sevcik KM, Johnson AT, Oetting TA, Haugsdal JM, et al. Metabolic and proteomic indications of diabetes progression in human aqueous humor. PloS one. 2023;18(1):e0280491.

61. Gao M, Monian P, Quadri N, Ramasamy R, Jiang X. Glutaminolysis and Transferrin Regulate Ferroptosis. Molecular Cell. 2015;59(2):298–308.

62. Jurkunas UV. Fuchs Endothelial Corneal Dystrophy Through the Prism of Oxidative Stress. Cornea. 2018;37 Suppl 1:S50–s4.

63. Liu J, Kuang F, Kroemer G, Klionsky DJ, Kang R, Tang D. Autophagy-Dependent Ferroptosis: Machinery and Regulation. Cell Chem Biol. 2020;27(4):420–35.

64. Levi S, Corsi B, Bosisio M, Invernizzi R, Volz A, Sanford D, et al. A Human Mitochondrial Ferritin Encoded by an Intronless Gene. J Biol Chem. 2001;276(27):24437–40.

65. Yang H, Yang M, Guan H, Liu Z, Zhao S, Takeuchi S, et al. Mitochondrial ferritin in neurodegenerative diseases. Neurosci Res. 2013;77(1-2):1–7.

66. Gao G, Chang Y-Z. Mitochondrial ferritin in the regulation of brain iron homeostasis and neurodegenerative diseases. Front Pharmacol. 2014;5(19).

67. Sui X, Zhang R, Liu S, Duan T, Zhai L, Zhang M, et al. RSL3 Drives Ferroptosis Through GPX4 Inactivation and ROS Production in Colorectal Cancer. Front Pharmacol. 2018;9:1371-.

68. Bai Y, Meng L, Han L, Jia Y, Zhao Y, Gao H, et al. Lipid storage and lipophagy regulates ferroptosis. Biochemical and Biophysical Research Communications. 2019;508(4):997–1003.

69. Riegman M, Sagie L, Galed C, Levin T, Steinberg N, Dixon SJ, et al. Ferroptosis occurs through an osmotic mechanism and propagates independently of cell rupture. Nat Cell Biol. 2020;22(9):1042–8.

70. Tyrrell RM. Activation of mammalian gene expression by the UV component of sunlight--from models to reality. Bioessays. 1996;18(2):139–48.

71. Pourzand C, Watkin RD, Brown JE, Tyrrell RM. Ultraviolet A radiation induces immediate release of iron in human primary skin fibroblasts: the role of ferritin. Proc Natl Acad Sci U S A. 1999;96(12):6751–6.

72. Valerio HP, Ravagnani FG, Yaya Candela AP, Dias Carvalho da Costa B, Ronsein GE, Di Mascio P. Spatial proteomics reveals subcellular reorganization in human keratinocytes exposed to UVA light. iScience. 2022;25(4):104093.

73. Jugé R, Breugnot J, Da Silva C, Bordes S, Closs B, Aouacheria A. Quantification and Characterization of UVB-Induced Mitochondrial Fragmentation in Normal Primary Human Keratinocytes. Scientific Reports. 2016;6(1):35065.

74. Bersuker K, Hendricks JM, Li Z, Magtanong L, Ford B, Tang PH, et al. The CoQ oxidoreductase FSP1 acts parallel to GPX4 to inhibit ferroptosis. Nature. 2019;575(7784):688-92.

75. Fan Z, Wirth AK, Chen D, Wruck CJ, Rauh M, Buchfelder M, et al. Nrf2-Keap1 pathway promotes cell proliferation and diminishes ferroptosis. Oncogenesis. 2017;6(8):e371.

76. Hou W, Xie Y, Song X, Sun X, Lotze MT, Zeh HJ, et al. Autophagy promotes ferroptosis by degradation of ferritin. Autophagy. 2016;12(8):1425–8.

77. Xu C, Sun S, Johnson T, Qi R, Zhang S, Zhang J, et al. The glutathione peroxidase Gpx4 prevents lipid peroxidation and ferroptosis to sustain Treg cell activation and suppression of antitumor immunity. Cell Rep. 2021;35(11):109235.

78. Geng N, Shi BJ, Li SL, Zhong ZY, Li YC, Xua WL, et al. Knockdown of ferroportin accelerates erastin-induced ferroptosis in neuroblastoma cells. Eur Rev Med Pharmacol Sci. 2018;22(12):3826–36.

79. Ward DM, Kaplan J. Ferroportin-mediated iron transport: expression and regulation. Biochimica et biophysica acta. 2012;1823(9):1426–33.

80. Harada N, Kanayama M, Maruyama A, Yoshida A, Tazumi K, Hosoya T, et al. Nrf2 regulates ferroportin 1-mediated iron efflux and counteracts lipopolysaccharide-induced ferroportin 1 mRNA suppression in macrophages. Archives of biochemistry and biophysics. 2011;508(1):101–9.

81. Mao C, Liu X, Zhang Y, Lei G, Yan Y, Lee H, et al. DHODH-mediated ferroptosis defence is a targetable vulnerability in cancer. Nature. 2021;593(7860):586-90.

82. Gain P, Jullienne R, He Z, Aldossary M, Acquart S, Cognasse F, et al. Global Survey of Corneal Transplantation and Eye Banking. JAMA Ophthalmol. 2016;134(2):167–73.

83. Vile GF, Tyrrell RM. UVA radiation-induced oxidative damage to lipids and proteins in vitro and in human skin fibroblasts is dependent on iron and singlet oxygen. Free radical biology & medicine. 1995;18(4):721–30.

84. Morlière P, Moysan A, Santus R, Hüppe G, Mazière JC, Dubertret L. UVA-induced lipid peroxidation in cultured human fibroblasts. Biochim Biophys Acta. 1991;1084(3):261–8.

85. Punnonen K, Jansén CT, Puntala A, Ahotupa M. Effects of in vitro UVA irradiation and PUVA treatment on membrane fatty acids and activities of antioxidant enzymes in human keratinocytes. The Journal of investigative dermatology. 1991;96(2):255–9.

86. Chu Y, Hu J, Liang H, Kanchwala M, Xing C, Beebe W, et al. Analyzing pre-symptomatic tissue to gain insights into the molecular and mechanistic origins of late-onset degenerative trinucleotide repeat disease. Nucleic acids research. 2020;48(12):6740–58.

87. Dobin A, Davis CA, Schlesinger F, Drenkow J, Zaleski C, Jha S, et al. STAR: ultrafast universal RNA-seq aligner. Bioinformatics. 2013;29(1):15–21.

88. Robinson MD, McCarthy DJ, Smyth GK. edgeR: a Bioconductor package for differential expression analysis of digital gene expression data. Bioinformatics. 2010;26(1):139–40.

89. Wisniewski JR, Zougman A, Nagaraj N, Mann M. Universal sample preparation method for proteome analysis. Nat Methods. 2009;6(5):359–62.

90. Rappsilber J, Mann M, Ishihama Y. Protocol for micro-purification, enrichment, pre-fractionation and storage of peptides for proteomics using StageTips. Nat Protoc. 2007;2(8):1896–906.

91. Chambers MC, Maclean B, Burke R, Amodei D, Ruderman DL, Neumann S, et al. A cross-platform toolkit for mass spectrometry and proteomics. Nat Biotechnol. 2012;30(10):918–20.

92. Sturm M, Bertsch A, Gropl C, Hildebrandt A, Hussong R, Lange E, et al. OpenMS - an open-source software framework for mass spectrometry. BMC Bioinformatics. 2008;9:163.

93. Craig R, Beavis RC. TANDEM: matching proteins with tandem mass spectra. Bioinformatics. 2004;20(9):1466–7.

94. Geer LY, Markey SP, Kowalak JA, Wagner L, Xu M, Maynard DM, et al. Open mass spectrometry search algorithm. J Proteome Res. 2004;3(5):958–64.

