## Supplementary material for "*TCF4* trinucleotide repeat expansions and UV irradiation increase susceptibility to ferroptosis in Fuchs endothelial corneal dystrophy"

**Supplementary figure 1: 4-HNE expression in FECD surgical and healthy control tissues.** Left, black and white figure represents antibody detection and right, blue colored figure represents total protein. Most dominant protein modification bands were selected, quantified and added together.

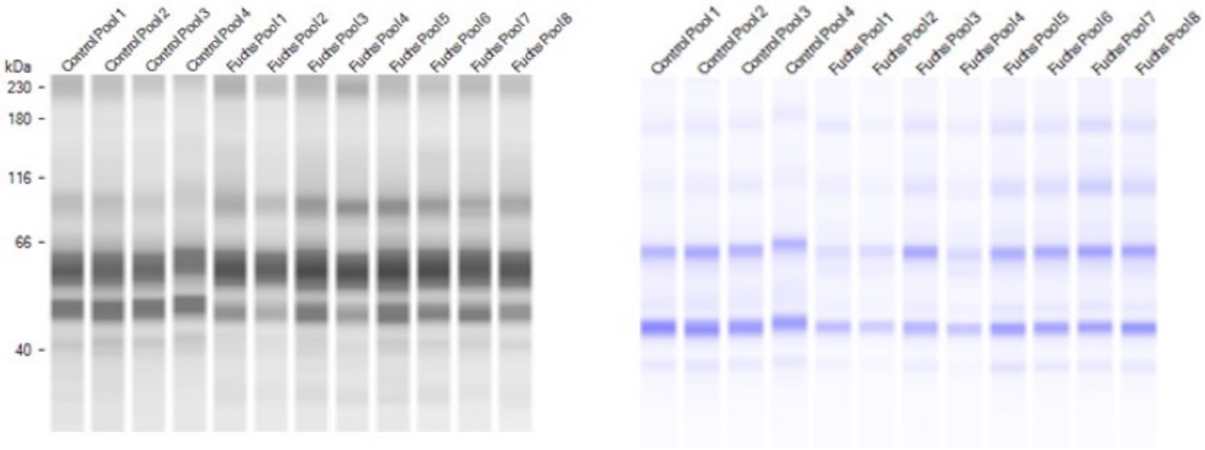

**Wave-like ferroptosis mediated death pattern in FECD cells:**

F35T cells were seeded at 100,000 cells/well for 18 h. Cells were treated with 1 µM of RSL3 and 50 nM of live cell impermeable nuclear staining dye, SYTOX Green. Cells were imaged by confocal microscopy in regular frequency to observe the presence of green fluorescence coming from SYTOX green when cell dyes due to RSL3-induced ferroptosis. Multiple images were taken from each well to observe wave-like death pattern in F35T cells.

**Supplementary figure 2: Ferroptosis in F35T cells happens in wave-like pattern.** Images were captured from single well (right to left) after 5 h, where all the cells had same treatment. Majority of the cells on right side of the well were found to be dead detected by SYTOX Green, whereas on the left side, very few cells were dead in the same well. Measurement bars in the images are 400 µm.

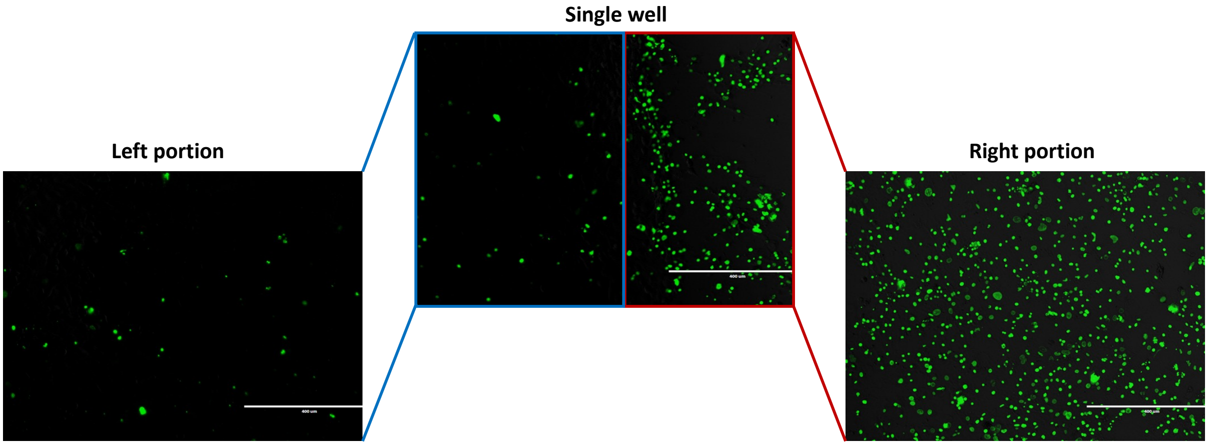

**UVA irradiation dose screening:**

UVA irradiation was tested for toxicity at different fluence levels between 5 and 25 J/cm$^2$ at 5 J/cm$^2$ increments. HCEC-F35T cells were seeded at 20,000 cells/well of 96 well plates and incubated for 18 h at 37°C and 5% $CO_2$. After incubation, cells were irradiated with different fluence levels of 5, 10, 15, 20, and 25 J/cm$^2$. Following UV irradiation, cell viability was measured using a MTS assay kit.

**Supplementary figure 3: Safe dose of UVA irradiation screening on HCEC-F35T cells.**

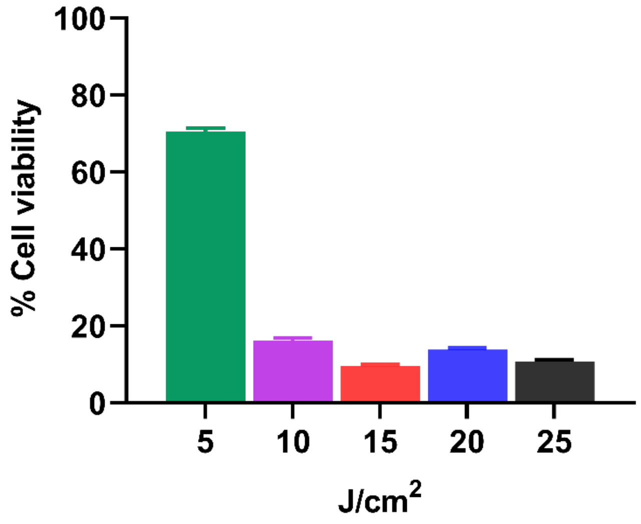

**TCF4 genotyping by Southern analysis:**

Genomic DNA was extracted from control and FECD immortalized cell lines using DNeasy Blood & Tissue Kit (Qiagen, Germantown, MD). Southern analysis was conducted by Lofstrand Labs (Gaithersburg, MD).

**Supplementary figure 4: Southern blot of genomic DNA samples showing *TCF4* intronic repeat allele sizes from a normal control corneal endothelial cell line (lane 1) and 2 FECD patient corneal endothelial cell lines (lanes 2 and 3).** *TCF4* repeat length genotyping was performed on genomic DNA extracted from immortalized cell lines. Lane 1 shows the control cell line HCEnT-21T (kindly provided by Ula Jurkunas, MD, Harvard Medical School) (1), and lane 2 shows an FECD cell line, both with repeat alleles in the normal length range. Lane 3 shows the FECD cell line F35T which has one normal and one expanded repeat length allele with approximately 4500 trinucleotide repeats.

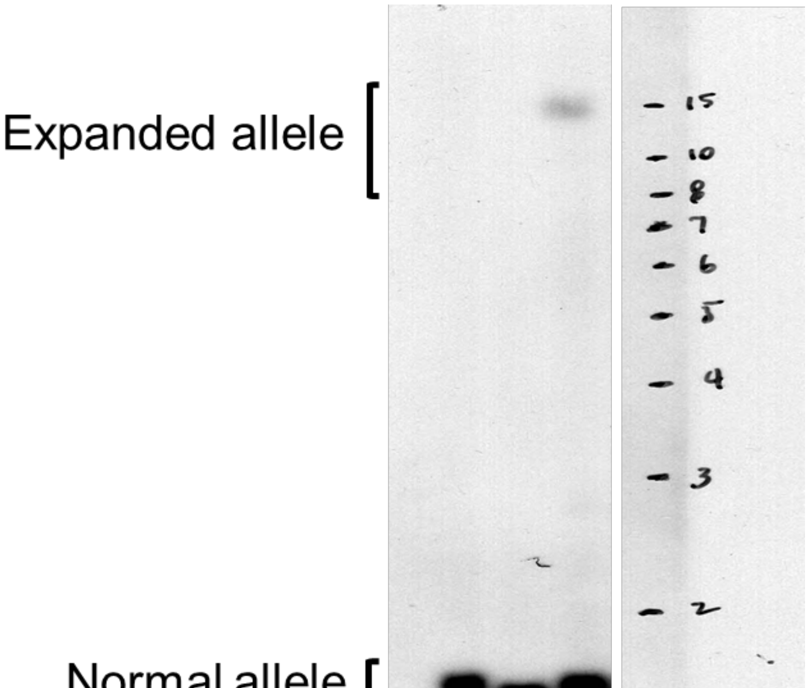

**Supplementary table 1: Demographics of donor EDM sample used in qPCR assays.**

| # | Diagnosis | Age | Sex | Medical history |
|---|---|---|---|---|
| 1 | Non FECD | 70 | M | Alcoholic cirrhosis, Gout, Hypothyroidism, HLD, Portal HTN, Small bowel and descending enterocolitis, Splenomegaly, Ascites, AKI, Umbilical hernia, Abdominal pain w/ nausea and vomiting, Allergy to Sulfa, Current every day smoker/tobacco use, EtOH use, Bilateral cataract extraction w/ IOL placements, Back Sx, Rotator cuff repair, Vasectomy, Paracentesis |
| 2 | Non FECD | 47 | M | Alcoholic cirrhosis, EtOH use, hemorrhagic shock d/t esophageal varices, tobacco use, decompensated liver disease, hyponatremia, secondary thrombocytopenia, acute blood loss anemia, banding sx. |
| 3 | Non FECD | 83 | F | Urinary incontinence, arthritis, glasses, blood transfusion, motion sickness, at risk for malignant hyperthermia, sleep apnea, high cholesterol, constipation, right knee pain, difficult intubation, h/o postoperative delirium, postoperative nausea and vomiting, pseudocholinesterase deficiency, spinal headache, hysterectomy, trigger finger release, carpal tunnel release, colonoscopy, cystoscopy, tubal ligation, bladder repair, joint replacement, breast biopsy, right total knee, cardiac catherization, coronary artery bypass, left atrial appendage ligation. |
| 4 | Non FECD | 90 | F | Allergic rhinitis, GERD, HTN, HLD, mitral valve prolapse, obesity, TIA, wide-angle glaucoma, cataract, cholecystectomy, hip replacement, knee replacement, wide-angle glaucoma, cataract, latanoprost ophthalmic. |
| 5 | Non FECD | 87 | M | COPD on $O_2$, chronic hypercapnic respiratory failure, cardiomyopathy, aortic stenosis, atrial flutter w/ablation, right heart failure, common variable immunodeficiency, bladder cancer, GERD, hypothyroidism, rheumatic fever, CABG, bioprosthetic aortic valve replacement. |
| 6 | Non FECD | 68 | F | esophageal varices w/banding, cirrhosis, radiation therapy, falls, colonic ulcers, endometrial cancer, GI bleed, laser therapy. |
| 7 | Non FECD | 66 | M | Dermatitis. |
| 8 | Non FECD | 67 | M | Alcohol use disorder, HTN, PTSD, Seizure, ADHD, Chronic lower extremity venous insufficiency, Peripheral neuropathy, Nonadherent w/ medications, Former tobacco use; Hypotensive presumed 2/2 septic shock, AKI, leukocytosis and anemia, Foley placement via cystoscopy, Arterial line placement. |
| 9 | Non FECD | 65 | M | Stage 4 esophageal cancer, chemoradiation, EtOH use, prior tobacco use, actinic keratosis, anxiety, depression, headache, HLD, HTN, varicella, basal cell carcinoma, gastrostomy placement, inguinal hernia repair, liver biopsy, flap reconstruction w/Mohs, septoplasty, bilateral cataract extraction. |
| 10 | Non FECD | 39 | M | Alcohol abuse, alcoholic liver cirrhosis, esophageal varices, tobacco use. |
| 11 | Non FECD | 42 | M | Alcohol abuse, cirrhosis. |
| 12 | Non FECD | 80 | M | Sigmoid colon carcinoma, monoclonal gammopathy, arthritis, HTN, chemotherapy, decubitus ulcers, malignant neoplasm of large intestine, prior tobacco use, EtOH use, chronic bilateral leg |

| | | | | |
|---|---|---|---|---|
| | | | | cellulitis, a-fib, colon sx, colonoscopy, partial colectomy w/coloproctostomy. |
| 13 | FECD | 95 | F | HTN, Hyperlipidemia, h/o bradycardia, atrial tachycardia, GERD, Benign neoplasm of the thyroid glands, Vit D deficiency, chronic kidney disease, osteoarthritis, ex-tobacco use, EtOH use. |
| 14 | FECD | 63 | F | Obesity, allergies, osteoarthrosis, EtOH use, appy, C-section. |
| 15 | FECD | 69 | M | Arthritis. |
| 16 | FECD | 79 | F | Hypothyroidism, osteopenia, hyperlipidemia, anxiety, ex-tobacco use. |
| 17 | FECD | 76 | F | Sleep apnea, HTN, hyperlipidemia, CAD, CHF, depression, anxiety, GERD, EtOH use. |
| 18 | FECD | 71 | M | Allergies, HTN, hyperlipidemia, paresthesias, GERD, pyogenic liver abscess, colonic diverticular abscess, cholecystectomy, ex-tobacco use, h/o alcoholism. |
| 19 | FECD | 78 | F | Breast cancer s/p resection, sleep apnea, h/o A-fib, GERD, breast ptosis, ankle sx, BSO, hysterectomy, lumpectomy of both breast. |
| 20 | FECD | 67 | F | HTN, hyperlipidemia, IBS, migraines, palpitations, GERD. |
| 21 | FECD | 77 | M | Thyroid disease. |
| 22 | FECD | 76 | F | Hypothyroidism, sensorineural hearing loss (SNHL) of both ears. |
| 23 | FECD | 73 | F | Arthritis, back problems s/p back fusion x2, allergies, tonsillectomy, hysterectomy, appy. |
| 24 | FECD | 76 | M | Hyperlipidemia, GERD, Vit D deficiency, eczema, osteoarthritis. |
| 25 | FECD | 70 | M | HTN, obesity, diastolic dysfunction, ventricular bigeminy, hyperlipidemia, osteoarthritis. |
| 26 | FECD | 77 | F | Allergic rhinitis, diverticulosis of colon, GERD, obesity, HTN, arrhythmia, carpal tunnel syndrome, appendectomy, carpal tunnel sx. |
| 27 | FECD | 65 | F | Hypothyroidism, A-fib, h/o malignant neoplasm of breast s/p mastectomy. |
| 28 | FECD | 78 | F | Hypothyroidism, breast augmentation, colectomy. |
| 29 | FECD | 71 | F | h/o PE, HTN, anxiety, GERD, spinal fusion, ex-tobacco use, EtOH use. |
| 30 | FECD | 63 | M | Inguinal hernia repair, tobacco use. |
| 31 | FECD | 75 | M | Hyperlipidemia, ex-tobacco use. |
| 32 | FECD | 63 | F | Sleep apnea, obesity, depression, GERD, breast cancer s/p resection, renal cell carcinoma, hysterectomy, EtOH use, h/o Tram flap reconstruction. |
| 33 | FECD | 80 | F | HTN, Hyperlipidemia, chronic lower back pain, h/o Meniere's disease, CVA, paresthesia, depression, hypothyroidism. |
| 34 | FECD | 67 | M | CAD s/p CABG, benign essential HTN, hyperlipidemia, benign prostatic hyperplasia, ex-tobacco use. |
| 35 | FECD | 75 | F | Hypercholesterolemia, osteoarthritis. |
| 36 | FECD | 72 | F | HTN, anxiety/nervousness, hyperlipidemia, obesity. |

Abbreviations: A-fib, atrial fibrillation; ASCVD, atherosclerotic cardiovascular disease; BiPAP, bilevel positive airway pressure; CAD, coronary artery disease; COPD, chronic obstructive pulmonary disease; CPAP, continuous positive airway pressure; DVT, deep vein thrombosis; EtOH, ethanol; HLD, hypersensitivity lung disease; HTN, hypertension; MI, myocardial infarction; NIDDM, non-insulin-dependent diabetes mellitus; OSA, obstructive sleep apnea; PE, pulmonary embolism; SOB, shortness of breath; w/, with; w/o, without.

**Supplementary table 2: Demographics of donor EDM sample used in western blot assays.**

| # | Diagnosis | Age | Sex | Medical history |
|---|---|---|---|---|
| 1 | Non FECD | 70 | M | Alcoholic cirrhosis, Gout, Hypothyroidism, HLD, Portal HTN, Small bowel and descending enterocolitis, Splenomegaly, Ascites, AKI, Umbilical hernia, Abdominal pain w/ nausea and vomiting, Allergy to Sulfa, Current every day smoker/tobacco use, EtOH use, Bilateral cataract extraction w/ IOL placements, Back Sx, Rotator cuff repair, Vasectomy, Paracentesis |
| 2 | Non FECD | 47 | M | Alcoholic cirrhosis, EtOH use, hemorrhagic shock d/t esophageal varices, tobacco use, decompensated liver disease, hyponatremia, secondary thrombocytopenia, acute blood loss anemia, banding sx. |
| 3 | Non FECD | 83 | F | Urinary incontinence, arthritis, glasses, blood transfusion, motion sickness, at risk for malignant hyperthermia, sleep apnea, high cholesterol, constipation, right knee pain, difficult intubation, h/o postoperative delirium, postoperative nausea and vomiting, pseudocholinesterase deficiency, spinal headache, hysterectomy, trigger finger release, carpal tunnel release, colonoscopy, cystoscopy, tubal ligation, bladder repair, joint replacement, breast biopsy, right total knee, cardiac catherization, coronary artery bypass, left atrial appendage ligation. |
| 4 | Non FECD | 90 | F | Allergic rhinitis, GERD, HTN, HLD, mitral valve prolapse, obesity, TIA, wide-angle glaucoma, cataract, cholecystectomy, hip replacement, knee replacement, wide-angle glaucoma, cataract, latanoprost ophthalmic. |
| 5 | Non FECD | 87 | M | COPD on $O_2$, chronic hypercapnic respiratory failure, cardiomyopathy, aortic stenosis, atrial flutter w/ablation, right heart failure, common variable immunodeficiency, bladder cancer, GERD, hypothyroidism, rheumatic fever, CABG, bioprosthetic aortic valve replacement. |
| 6 | Non FECD | 68 | F | esophageal varices w/banding, cirrhosis, radiation therapy, falls, colonic ulcers, endometrial cancer, GI bleed, laser therapy. |
| 7 | Non FECD | 66 | M | Dermatitis. |
| 8 | Non FECD | 67 | M | Alcohol use disorder, HTN, PTSD, Seizure, ADHD, Chronic lower extremity venous insufficiency, Peripheral neuropathy, Nonadherent w/ medications, Former tobacco use; Hypotensive presumed 2/2 septic shock, AKI, leukocytosis and anemia, Foley placement via cystoscopy, Arterial line placement. |
| 9 | Non FECD | 65 | M | Stage 4 esophageal cancer, chemoradiation, EtOH use, prior tobacco use, actinic keratosis, anxiety, depression, headache, HLD, HTN, varicella, basal cell carcinoma, gastrostomy placement, inguinal hernia repair, liver biopsy, flap reconstruction w/Mohs, septoplasty, bilateral cataract extraction. |
| 10 | Non FECD | 39 | M | Alcohol abuse, alcoholic liver cirrhosis, esophageal varices, tobacco use. |
| 11 | Non FECD | 42 | M | Alcohol abuse, cirrhosis. |
| 12 | Non FECD | 80 | M | Sigmoid colon carcinoma, monoclonal gammopathy, arthritis, HTN, chemotherapy, decubitus ulcers, malignant neoplasm of large intestine, prior tobacco use, EtOH use, chronic bilateral leg |

| | | | | |
|---|---|---|---|---|
| | | | | cellulitis, a-fib, colon sx, colonoscopy, partial colectomy w/coloproctostomy. |
| 13 | FECD | 78 | F | Pulmonary embolism, long term current use of anticoagulant therapy, mild pulmonary hypertension, GERD, hiatal hernia, temporal arteritis, current chronic use of systemic steroids, HTN, rotator cuff surgery, ex-tobacco use, EtOH use. |
| 14 | FECD | 56 | F | Choledocholithiasis, depression, S/P cholecystectomy, appy, C-section, hysterectomy, exploratory laparotomy, EtOH use. |
| 15 | FECD | 75 | M | Hyperlipidemia, ex-tobacco use. |
| 16 | FECD | 76 | F | Sleep apnea, HTN, hyperlipidemia, CAD, CHF, depression, anxiety, GERD, EtOH use. |
| 17 | FECD | 62 | M | GERD, diverticulosis, allergies. |
| 18 | FECD | 70 | F | Asthma, hyperlipidemia, obesity, anxiety/nervousness, GERD, hypothyroidism, ex-tobacco use. |
| 19 | FECD | 75 | F | Sleep apnea, HTN, CAD, obesity, CVA, Appy, cardiac stents, hysterectomy, cholecystectomy, bilateral knee replacements, T&A, EtOH use. |
| 20 | FECD | 66 | M | Hyperlipidemia, GERD. |
| 21 | FECD | 64 | F | Asthma, hyperlipidemia, h/o of SVT and A-fib, GERD, thyroid disease. |
| 22 | FECD | 63 | F | Obesity, allergies, osteoarthrosis, EtOH use, appy, C-section. |
| 23 | FECD | 71 | M | Allergies, HTN, hyperlipidemia, paresthesias, GERD, pyogenic liver abscess, colonic diverticular abscess, cholecystectomy, ex-tobacco use, h/o alcoholism. |
| 24 | FECD | 72 | F | HTN, anxiety/nervousness, hyperlipidemia, obesity. |
| 25 | FECD | 84 | F | HTN, CAD, GERD, cervical spondylosis with myelopathy and radiculopathy. |
| 26 | FECD | 92 | M | Hyperlipidemia, leg swelling, bilateral HA, GERD, osteoarthrosis. |
| 27 | FECD | 65 | M | Rotator cuff repair, EtOH use. |
| 28 | FECD | 75 | F | Asthma, chronic bronchitis and sinusitis, HTN, cardiac arrhythmia, GERD, h/o colon cancer s/p resection, sinus sx, endometrial ablation, tubal ligation, social EtOH use. |
| 29 | FECD | 80 | M | Dupuytren’s contracture, arthritis, ex-Tobacco use, EtOH use. |
| 30 | FECD | 60 | F | Asthma, allergies, obstructive sleep apnea, morbid obesity, iron (Fe) deficiency anemia, GERD, eczema, osteoarthritis, hyperlipidemia. |
| 31 | FECD | 67 | M | Mitral valve prolapse, left superior oblique palsy, testicular cancer s/p radiation therapy and resection, hernia repair. |
| 32 | FECD | 77 | F | Allergic rhinitis, diverticulosis of colon, GERD, obesity, HTN, arrhythmia, carpal tunnel syndrome, appendectomy, carpal tunnel sx. |
| 33 | FECD | 64 | F | Sleep apnea, obesity, depression, GERD, breast cancer s/p resection, renal cell carcinoma, hysterectomy, EtOH use, h/o Tram flap reconstruction. |
| 34 | FECD | 75 | M | CAD s/p PCI and RCA stent, HTN, hyperlipidemia, obesity, asthma, sick sinus syndrome, unstable angina, sleep apnea, GERD. |
| 35 | FECD | 67 | F | HTN, hyperlipidemia, IBS, migraines, palpitations, GERD. |

| | | | | |
|---|---|---|---|---|
| 36 | FECD | 62 | F | Asthma, migraines, depression, humerus fracture, fibroid (uterus), cervical dysplasia, osteoarthritis s/p total hip replacement, hyperlipidemia, breast lumpectomy, cholecystectomy, ex-tobacco use. |

Abbreviations: A-fib, atrial fibrillation; ASCVD, atherosclerotic cardiovascular disease; BiPAP, bilevel positive airway pressure; CAD, coronary artery disease; COPD, chronic obstructive pulmonary disease; CPAP, continuous positive airway pressure; DVT, deep vein thrombosis; EtOH, ethanol; HLD, hypersensitivity lung disease; HTN, hypertension; MI, myocardial infarction; NIDDM, non-insulin-dependent diabetes mellitus; OSA, obstructive sleep apnea; PE, pulmonary embolism; SOB, shortness of breath; w/, with; w/o, without.

**Supplementary table 3: Demographics of donor EDM sample used in cytosolic iron detection.**

| # | Diagnosis | Age | Sex | Medical history |
|---|---|---|---|---|
| 1 | Non FECD | 71 | M | Cyst removed from scrotum, tobacco use and EtOH use |
| 2 | Non FECD | 72 | M | Appendectomy, cardiovascular left circumflex stent placed, right fifth finger amputation, right Carpal tunnel syndrome, vasectomy, fatty tumor removal, atherosclerotic coronary disease, migraines, chronic tobacco use, chronic alcohol abuse, severe allergic rhinitis, SOB, wheezing, MI, end stage COPD, emphysema, HTN, HLD, anemia, hyponatremia, depression, trouble breathing with seasonal allergies, lower extremity issues of spider veins and cold feet. |
| 3 | Non FECD | 40 | M | Alcohol abuse, obesity, tobacco dependence, withdrawal seizures, seizures, smoked marijuana, snorted cocaine borderline cirrhosis, detox of substance abuse, appendectomy, and knee surgery. |
| 4 | Non FECD | 52 | M | Acid reflux, Barrett esophagus, blood clotting tendency, gastrointestinal disorder, HTN, long term use of anticoagulants, PE, venous thrombosis and embolism, sleep apnea- wears CPAP, bipolar disorder, depression, headaches, mental decline with stress, Factor V Leiden mutation- heterozygous, benign brain tumor, bilateral rotator cuff tears, rare EtOH use, kidney stones, hiatal hernia, esophageal issues, bilateral knee, brain tumor excision, blood clot removed and filter placed in right groin, gastric bypass, esophageal stretching, brain radiation and Greenfield filter. |
| 5 | Non FECD | 33 | M | Suicide attempt by overdose, symptoms of suicidality and behavior changes, convulsions, poison ivy, seizures, refractory epilepsy, mouth sore, personality changes, and multiple seizures and occasional EtOH use. |
| 6 | Non FECD | 70 | M | NIDDM type 2 w/o complication, primary HTN, dyslipidemia, multiple MI, benign prostatic hyperplasia w/o lower urinary tract symptoms, CPAP dependence, nonalcoholic fatty live disease, OSA on BiPAP, ASCVD, paroxysmal A-fib w/ aberrancy, wide-complex tachycardia, paroxysmal tachycardia, hypertrophy of prostate w/o urinary obstruction and other lower urinary tract symptoms, large cecal polyp, multiple colonic polyps, CAD-multivessel, mild left ventricular dysfunction w/o evidence of volume overload, postoperative nausea and vomiting, stable angina, morbid obesity, bilateral lower extremity varicose veins, unspecified heart issues, former chronic tobacco use, rare EtOH use, cardiac catheterization w/ stents placed, right carpal tunnel release and laparoscopic cholecystectomy. |
| 7 | Non FECD | 36 | M | Retaining water, 3 teeth removed, ankle surgery, seasonal allergies, snorted/smoked meth, heroin use, smoked marijuana, ingested acid, ingested shrooms, tobacco use, COPD, CHF, ETOH, HTN, snoring, acute respiratory failure w hypoxia, morbid obesity, anxiety, thrombophilia, iron deficiency, |

| | | | | |
|---|---|---|---|---|
| | | | | insomnia, GERD, burn of right hand, drug eruption, discitis of thoracic region, bilateral lower extremity edema, salmonella bacteremia, substance abuse. |
| 8 | FECD | 88 | F | Cardiovascular disease, DVT, essential hypertension, pain in joint and shoulder region, disorders of bursae and tendons in shoulder region, diffuse cystic mastopathy, abnormal mammogram and history of stroke. |
| 9 | FECD | 78 | F | Stiffness of joint |
| 10 | FECD | 74 | M | Oncology, malignant neoplasm of overlapping sites of bladder, cardiovascular, A-fib, history of heart bypass surgery, systolic heart failure, cornea replaced by transplant, age-related nuclear cataract, bilateral, history of retinal tear and hyponatremia. |
| 11 | FECD | 88 | M | Cardiovascular, atrial flutter, presence of cardiac pacemaker, DVT, primary hypertension, history of coronary artery bypass surgery, ventricular tachycardia, endocrine hypoactive thyroid, corneal transplant, genitourinary, benign prostatic hyperplasia without urinary obstruction, Injury of globe of eye and pseudophakia. |
| 12 | FECD | 68 | F | No history |
| 13 | FECD | 73 | F | Impaired glucose tolerance, vitamin D deficiency, history of cornea transplant, genitourinary, cystocele, urge incontinence, osteopenia, pseudophakia, lactase deficiency and post-menopausal. |
| 14 | FECD | 62 | F | Asthma, rheumatoid arthritis, migraines, bilateral flat feet, hammer toe, numbness left leg, GERD, rare EtOH use, FECD, previous hallux valgus reconstructive surgery in the left foot. |

Abbreviations: A-fib, atrial fibrillation; ASCVD, atherosclerotic cardiovascular disease; BiPAP, bilevel positive airway pressure; CAD, coronary artery disease; COPD, chronic obstructive pulmonary disease; CPAP, continuous positive airway pressure; DVT, deep vein thrombosis; EtOH, ethanol; HLD, hypersensitivity lung disease; HTN, hypertension; MI, myocardial infarction; NIDDM, non-insulin-dependent diabetes mellitus; OSA, obstructive sleep apnea; PE, pulmonary embolism; SOB, shortness of breath; w/, with; w/o, without.

**Supplementary table 4: Demographics of donor EDM sample used in proteomics.**

| # | Diagnosis | Age | Sex | Other ocular diagnosis |
|---|---|---|---|---|
| 1 | Non FECD | 77 | F | Dry eye, MGD |
| 2 | Non FECD | 72 | F | No history |
| 3 | Non FECD | 82 | M | MGD |
| 4 | Non FECD | 80 | F | Amblyopia, exotropia |
| 5 | FECD | 64 | F | No history |
| 6 | FECD | 72 | M | MGD |
| 7 | FECD | 79 | F | No history |
| 8 | FECD | 66 | F | No history |

Abbreviations: FECD, Fuchs endothelial corneal dystrophy; MGD, meibomian gland dysfunction.

**Supplementary table 5**: FECD versus control aqueous humor (**A**) proteomics results and (**B**) ANOVA statistical analysis.

**Supplementary table 6**: Protein pathways significantly altered in FECD aqueous humor and corresponding proteins identified in FECD and control patient aqueous humor samples determined using IPA software.

**Supplementary table 7: Demographics of donor EDM sample used in cytosolic iron detection upon ultraviolet A exposure.**

| # | Diagnosis | Age | Sex | Medical history |
|---|---|---|---|---|
| 1 | Non FECD | 71 | F | Environmental allergies, blood transfusion w/o reported diagnosis, heart murmur, osteoporosis, HLD, osteopenia, ventricular septal defect s/p repair, seasonal hay fever and varicose veins. |
| 2 | Non FECD | 68 | M | Essential HTN, constant wheezing, chronic cough, significant coronary artery calcifications, clavicle fracture, fracture of left hip, fracture of lumbar spine T10-L2, glenoid fracture of shoulder, squamous cell carcinoma of skin, lung collapse, pleural effusion on right, pulmonary nodule, rib fractures, tibial fracture, scratch/bite by pet dog, trauma to back, shoulder/collar bone and leg due to fall, skin and lymph node cancer, radiation therapy, chemotherapy, chronic tobacco use, EtOH abuse and allergy to penicillin. |
| 3 | Non FECD | 60 | M | Seasonal allergies, smoked marijuana, previous tobacco use and EtOH use. |

Abbreviations: EtOH, ethanol; HLD, hypersensitivity lung disease; HTN, hypertension; w/o, without.

**Supplementary table 8: List of primers used in qPCR.**

| Gene | Primer sequences | Amplicon size (bp) |
|---|---|---|
| *FTL* | Forward: 5`-AGGCCCTTTTGGATCTTCAT-3`<br>Reverse: 5`-CAGGTGGTCACCCATCTTCT-3` | 122 |
| *FTH* | Forward: 5`-CTGAATGCAATGGAGTGTGC-3`<br>Reverse: 5`-AATGGGGGTCATTTTTGTCA-3` | 97 |
| *FSP1* | Forward: 5`-GGCCAACATCGTCAACTCTG-3`<br>Reverse: 5`-ACACCGTCATTTCTCCCCAT-5` | 96 |
| *GPX4* | Forward: 5`-TCAGCAAGATCTGCGTGAAC-3`<br>Reverse: 5`-GGGGCAGGTCCTTCTCTATC-3` | 195 |
| *SLC40A1 (FPN1)* | Forward: 5`-CAGGGACTGAGTGGTTCCAT-3`<br>Reverse: 5`-ACCACATTTTCGACGTAGCC-3` | 102 |
| *TFR1* | Forward: 5`-ACCATTGTCATATACCCGGTTCA-3`<br>Reverse: 5`-CAATAGCCCAAGTAGCCAATCAT-3` | 219 |

**Supplementary video 1:** Time-lapse video imaging of ferroptosis in FECD induced by RSL3 with fluorophores visualizing movement of lipid droplets by LipidSpot™ 610 and terminal cell death by SYTOX green. **IA:** Ferroptosis in B4G12 cells; **1B:** Ferroptosis in F35T cells.

**Supplementary video 2:** Time-lapse video imaging of ferroptosis in FECD induced by RSL3 showing cell morphology in brightfield confocal microscopy. **2A:** Ferroptosis in B4G12 cells; **2B:** Ferroptosis in F35T cells.

**Supplementary video 3:** Time-lapse video imaging of ferroptosis in FECD induced by 1.5 $J/cm^2$ UVA irradiation showing cell morphology in brightfield confocal microscopy and fluorophores visualizing movement of lipid droplets by LipidSpot™ 610 and terminal cell death by SYTOX green. **3A:** Ferroptosis in B4G12 cells; **3B:** Ferroptosis in F35T cells.

**References**

1. Schmedt T, Chen Y, Nguyen TT, Li S, Bonanno JA, Jurkunas UV. Telomerase immortalization of human corneal endothelial cells yields functional hexagonal monolayers. PloS one. 2012;7(12):e51427.
